# Get the gist of the story: Neural map of topic keywords in multi-speaker environment

**DOI:** 10.1101/2022.05.05.490770

**Authors:** Hyojin Park, Joachim Gross

## Abstract

Neural representation of lexico-semantics in speech processing has been revealed in recent years. However, to date, how the brain makes sense of the higher-level semantic gist (topic keywords) of a continuous speech remains mysterious. Capitalizing on a generative probabilistic topic modelling algorithm on speech materials to which participants listened while their brain activities were recorded by Magnetoencephalography (MEG), here we show spatio-temporal neural representation of topic keywords in a multi-speaker environment where task-relevant (attended) and -irrelevant (unattended) speech co-exits. We report the difference of neural representation between salient and less salient semantic gist of both attended and unattended speech. Moreover, we show that greater sensitivity to semantically salient unattended speech in the left auditory and motor cortices negatively mediates attended speech comprehension.

## Introduction

Humans have the remarkable ability to get the gist of the story. Imagine that you are listening to a talk without any prior information about the topic of the talk. As the talk unfolds, you will identify keywords that will enable you to infer the topic of the talk. How does the human brain extract topic keywords from the story, and how does the brain distinguish critical, relevant words (i.e., keywords) from less critical, irrelevant words? What are the spatio-temporal characteristics in brain activity that reflect the identification and processing of topic keywords, particularly in the context of effortful listening, for instance, at a cocktail party?

An increasing number of studies has demonstrated that neural representations of acoustic (Park et al., 2015; Park et al., 2016; Hauswald et al., 2018; Park et al., 2018b; Park et al., 2018a; Biau et al., 2021; Haider et al., 2022) and linguistic, e.g., semantic features (Kutas and Federmeier, 2011; Strauss et al., 2014; Huth et al., 2016; Wang et al., 2018; Broderick et al., 2019; Kaufeld et al., 2020) of naturalistic auditory or audiovisual speech are quantifiable in Magnetoencephalography (MEG) or Electroencephalography (EEG) recordings based on frequency-domain synchronization analysis or time-domain regression analysis. Furthermore, recent developments of natural language processing (NLP) models based on machine learning algorithms, such as word vectors (Mikolov et al., 2013), have brought breakthroughs not only to the area of artificial intelligence (AI), for example, speech/text recognition, machine translations (e.g., speech-to-text), but also to the neuroscientific study of rich, naturalistic speech stimuli (Broderick et al., 2018; Pereira et al., 2018) or movie (Nishida et al., 2021). For example, Broderick et al. (2018) has shown that using a trained computational language model, semantic vectors for content words of the continuous stimuli, which were given to participants during EEG recordings, can be used to identify semantic neural correlates.

Traditionally, neural semantic processing has been studied through semantic violations (Kutas and Federmeier, 2011). More recently, semantic analysis has been extended to word-level prediction (Wang et al., 2018) and even to the processing of continuous speech (Broderick et al., 2018; Broderick et al., 2019; Koskinen et al., 2020). However, the field still lacks the quantification of neural processes involved in higher-level semantic processing, such as investigating how the brain understands the topic of the story (main idea, keywords, semantic core). Comprehension of sentences in continuous speech requires multiple levels of hierarchical processing. One of the key components in this processing is the understanding of word meanings, i.e., lexico-semantic processing, which is a critical building block supporting the construction of a semantic core of the story. However, the understanding of latent meanings in idiomatic expressions, for example, cannot be attained through word-level processing. The comprehension rather builds upon complex contextual information. The neural mechanism underlying such processing remains unclear. As such, in the current study, we aim to investigate how the brain extracts the main topic in a continuous speech and how this process is implemented in multi-speaker environment where task-relevant (attended) and -irrelevant (unattended) stimuli co-exist.

To investigate brain responses engaging in the understanding of the main topic of the story, we first identified topic keywords in the spoken speech materials delivered to the participants during MEG measurement. Here we used a topic model algorithm, Latent Dirichlet Allocation (LDA), a text mining technique that has been developed to delineate short descriptions of the collection of text corpora (Blei et al., 2003) on speech chunks segmented in a perceptually relevant manner. LDA analysis results in sets of topic keywords and the probability of the words belonging to the topic. Thus, LDA enables identifying the topic probability for each speech chunk. For characterizing the brain representations of the topic keywords, the segmented speech chunks were sorted according to topic probability and divided into two conditions (high vs. low topic probability condition). Speech envelope in each condition and corresponding brain activities were used to fit encoding and decoding models of the multivariate temporal response function (mTRF).

Results from the decoding model prediction accuracy show that speech chunks with high topic probability are better reconstructed when compared to speech chunks with low topic probability within the attended and unattended talk. Strikingly, this was evident between speech chunks with high topic probability from the unattended talk and speech chunks with low topic probability from the attended talk. Moreover, we provide evidence that the processing of unattended speech chunks with semantic gist in the left frontal, auditory and motor areas negatively mediates attended speech comprehension. We show, for the first time to our knowledge, how the brain processes topic keywords in a continuous story in a challenging listening situation in a behaviourally relevant manner.

## Results

### Speech data processing and computational topic modelling

#### Segmentation of natural speech

Embedded linguistic units (e.g., embedded phrases or conjoined sentences) in natural speech and its temporal complexity with varying lengths of phrases and sentences have made it difficult to study high-level semantic processing beyond word-level during natural speech perception. Those time-varying semantic chunks are often marked by intermittent pauses (i.e., silences) from the speaker before moving to the next phrases or sentences. In order to derive topic keywords from a talk via a computational modelling algorithm, we first segmented each talk into phrases or sentences based on these acoustic properties. Using the library Syllable Nuclei (de Jong and Wempe, 2009) in Praat software (Boersma and Weenink, 2018), we obtained acoustic chunks with parameters of 0.25 s minimum pause (silence) durations, -25 dB silence threshold, and 1 s minimum duration of each speech chunk (Fig. 1a). We detected 129 speech chunks on average with a mean duration of 3.37 s from the seven talks used in the current study. We calculated speech rates by the number of syllables divided by the duration of the talk. For detecting peak intensity, 2 dB above the median was used as a threshold (the minimum dip between peaks). The profiles of the talks – for example, the number of syllables, speech rate, articulation rate (the number of syllables divided by phonation (speaking) time), the number of speech chunks for each talk – are reported in Supplementary Table 1.

**Figure 1.**
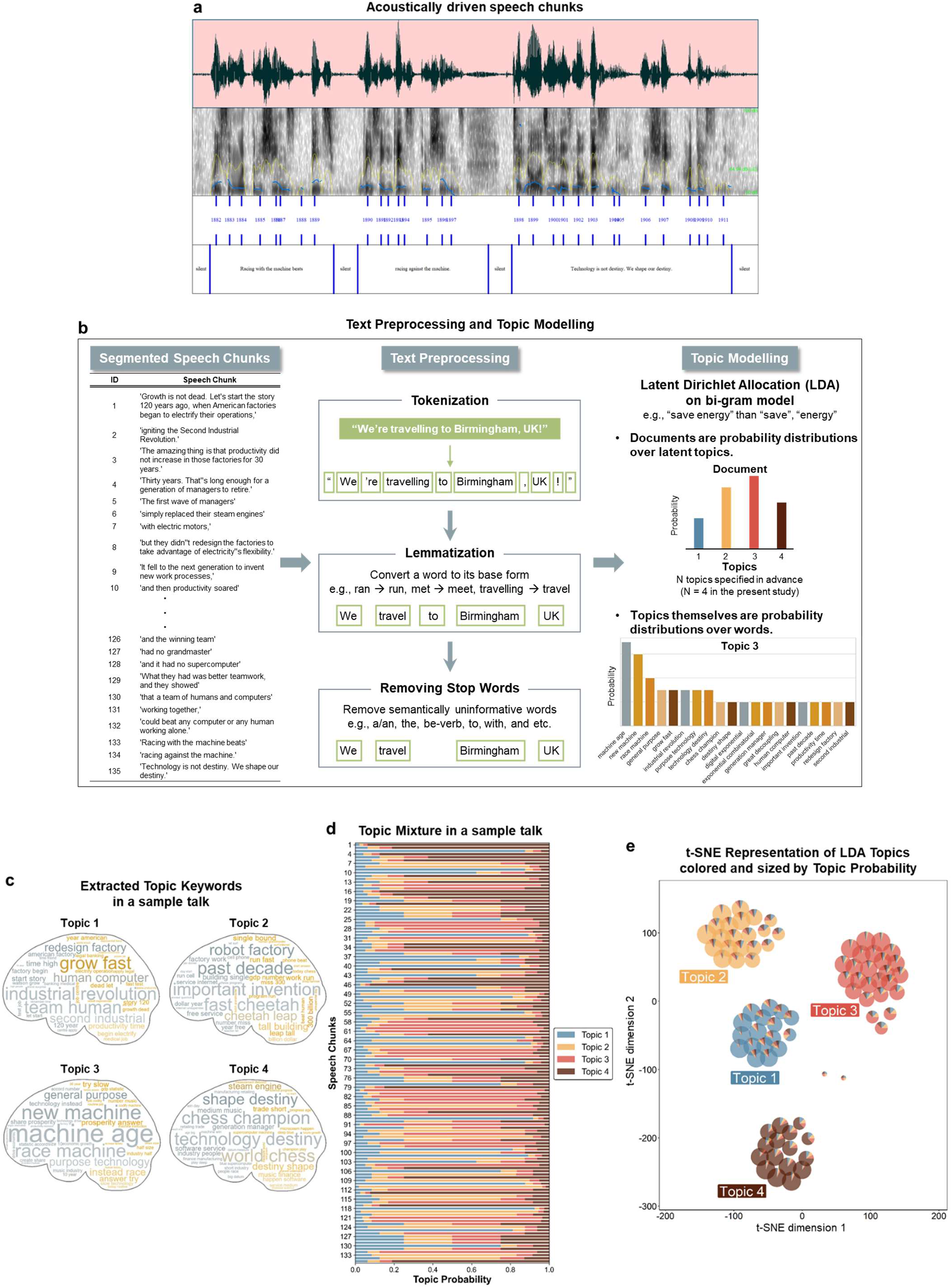
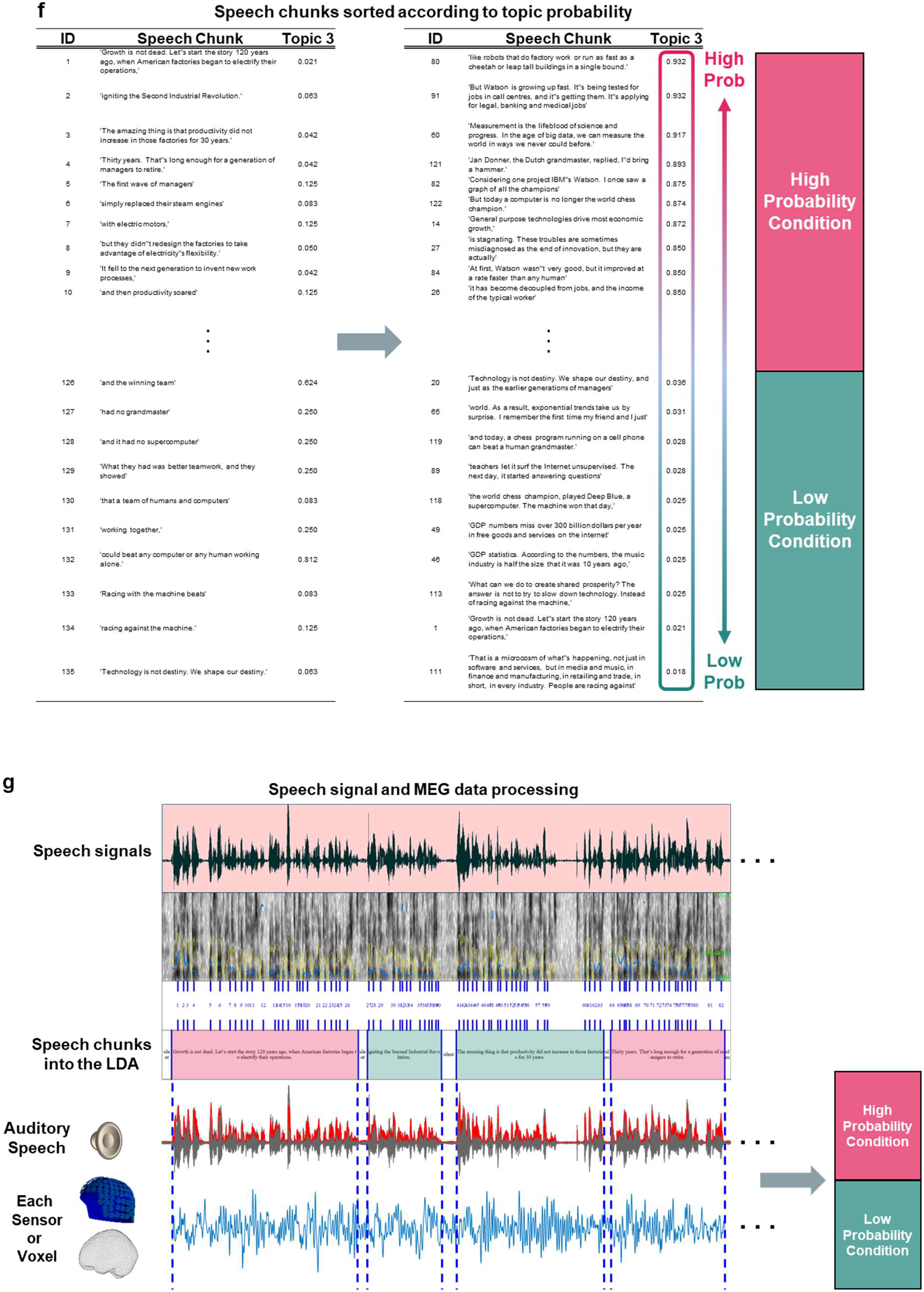
Segmentation of speech and topic modelling application. **a, Perceptually driven segmentation of speech**. A continuous speech was segmented into phrases or sentences in a perceptually relevant manner. Using the library Syllable Nuclei in Praat, we obtained acoustic speech chunks by the following thresholding parameters: pause (silence) duration (minimum 0.25 s long), loudness (−25 dB silence), and the length of speech chunks (minimum 1 s long). We detected 129 speech chunks on average with a mean duration of 3.37 s for seven continuous talks used in the study (also see Supplementary Table 1). A snapshot from one example talk is shown in the figure. Each row depicts a raw speech signal, spectrogram with intensity (in yellow) and pitch (in blue), the number of syllables, and annotations. **b, Text preprocessing and topic modelling**. Left column: An example of segmented speech chunks from a representative talk is shown. In this talk, 135 speech chunks were obtained. Middle column: Preprocessing of annotations of spoken speech materials was performed using a python library, spaCy, through the following three steps: tokenization, lemmatization and removal of stop words. Right column: Schematic illustration of topic modelling algorithm, Latent Dirichlet Allocation (LDA), is displayed. A fixed number of topics was specified in advance (4 in the current study), and bi-gram model was used in the topic model to optimally capture topic messages. **c, Extracted topic keywords**. Out of the LDA model, documents (referred to as speech chunks in the current study) assigned to topic *t* and words with high probability for topic *t* are obtained as outputs (document-topic matrix). The most common words with the highest probability are shown for each topic. **d, Distribution of topic probability across speech chunks in a representative talk**. Vectors of topic probabilities for each speech chunk were depicted using a stacked bar chart (topic mixture) where x- and y-axes depict topic probability and the identity of speech chunks, respectively. Color-coded bars in each row represent the probability distribution across 4 topics in a given speech chunk. In this sample talk, topic 3 has the highest probability across all speech chunks, which indicates the representative topic keywords for this talk. **e, t-SNE visualization of topics colored and sized by topic probability**. The document-topic matrix from the LDA model was subjected to t-SNE dimensionality reduction model, which was then fitted to be visualized in 2-d embedded space scatterpie chart. Each data point as a scatterpie chart represents each speech chunk which is clustered into a certain topic according to the highest probability for the topic. The scatterpie chart also shows the probability distribution over topics. **f, Speech chunks sorted according to the topic probability of the representative topic** (topic 3 in this example talk). The speech chunks shown in b were sorted from highest to lowest topic probability as to topic 3. To investigate the neural processing of topic representation, we split these speech chunks into high vs. low topic probability conditions. **g, Segmentation and allocation of the MEG and speech signals into topic probability conditions**. Corresponding brain data at sensor and source level and auditory speech envelope were split into high vs. low topic probability conditions as well, resulting in trial-based epochs.

#### Text preprocessing

Transcriptions of TED talks for video filmings were double-checked after the recordings by a professional speaker for a possible difference between transcriptions and actual speech during the filmings. Different parts were updated, and these finalized transcriptions were used for the annotations of the segmented speech chunks derived by the Syllable Nuclei (see the first column in Fig. 1b). Preprocessing of text materials was performed using a python library, spaCy (Honnibal and Montani, 2017), an open-source software library for advanced natural language processing (NLP), which is similar to NLTK (Natural Language Toolkit) – a popular open-source library released in 2001 (Bird et al., 2009). The preprocessing can be summarized into the following three steps (the middle column in Fig. 1b) – 1) tokenization, 2) lemmatization, and 3) removal of stop words, described below in more detail.

Using an English language model - OntoNotes 5 (Weischedel et al., 2013) by Linguistic Data Consortium (LDC), trained on a large corpus comprising various genres of text (news, conversational telephone speech, weblogs, usenet newsgroups, broadcast, talk shows) - embedded in spaCy, raw texts were tokenized into component pieces, e.g., prefix, suffix, infix and other exceptions. Next, these basic building blocks of document objects (tokens) were lemmatized, which is similar to stemming (word reduction, e.g., “boat” for “boats”, “boating”, “boater”) in other libraries, which works well, but with some issues given numerous exceptions in English (e.g., irregular verbs such as begin-began-begun). Lemmatization implements beyond the stemming (word reduction) and considers a language’s full vocabulary to apply a morphological analysis to words. For instance, the lemma of ‘was’ is ‘be’, and the lemma of ‘mice’ is ‘mouse’. In some cases, the lemma of ‘meeting’ might be ‘meet’ or ‘meeting’ depending on the context (e.g., “I’m meeting my boss tomorrow at the meeting.”). Lemmatization in spaCy is assumed to provide correct lemmas even in such cases by considering surrounding texts to determine a word’s part of speech, which has motivated our use of the spaCy library in the preprocessing steps for the subsequent topic model analysis. Then, stop words, which are presumed to be semantically uninformative in representing a content of a text, such as “a/an”, “the”, be-verbs, “and”, “with”, “seems”, “also” etc. were filtered out from the text. We used a ∼300 stop words list built in a machine learning python library, scikit-learn (Pedregosa et al., 2011) feature extraction module for texts (sklearn.feature_extraction.text). Furthermore, to rule out the possibility of emotional valence driven topic keywords representation in the brain, we confirmed that all the speech materials used in the study are with neutral sentiment by sentiment analysis (see Supplementary Table 2 for more details).

#### Topic modelling

Topic models are collective algorithms that try to uncover the hidden thematic structure in document collections (Blei, 2012). Recent advances in state-of-the-art machine learning algorithms and the era of big data enable the developments of the subfield of Natural Language Processing (NLP) at different levels, such as syntactic, lexical semantics, or discourse. The current study aims to map spatiotemporal neural representation of topic processing during natural speech perception, so we used one of the widely-used topic modelling algorithms, Latent Dirichlet Allocation (LDA) (Blei et al., 2003) in order to extract topic keywords in a talk. Latent Semantic Analysis (LSA) (Landauer and Dumais, 1997) is another widely used topic model (Hoffman, 2019). LDA and LSA are conceptually similar and allow us to efficiently analyze a large volume of text data by clustering documents into a certain number of topics. They are unsupervised learning algorithms as the text data is unlabelled. The assumption behind the topic modelling is that documents with similar topics use similar groups of words, so latent topics can be detected by the frequent co-occurrence of groups of words in documents across the corpus. Both LDA and LSA algorithms operate on word-document co-occurrence matrices in a low-dimensional space so that the occurrence of sets of words in multiple documents can be computed. LSA uses singular value decomposition (SVD) that the created singular vector of the co-occurrence assumed to be orthogonal. LDA is a generative probabilistic model as its name based off (Dirichlet distribution) that produces probabilities that a word derived from the distribution for a particular topic. More detailed differences between LSA and LDA are discussed in the previous literature (Griffiths et al., 2007; Pereira et al., 2011).

#### Latent Dirichlet Allocation (LDA) analysis

LDA topic model analysis was performed using the scikit-learn (Pedregosa et al., 2011) (class: sklearn.decomposition.LatentDirichletAllocation), following the preprocessing of texts described above. First, the preprocessed text documents are converted to a matrix of token counts (i.e., a sparse representation of token counts, also known as document-term matrix via scikit-learn class, sklearn.feature_extraction.text.CountVectorizer), then the LDA model is fitted to the document-term matrix.

LDA (Blei; Chen; Blei et al., 2003; Blei, 2012) is a generative probabilistic model for collections of texts implementing a three-level hierarchical Bayesian model. In the model, each item of a collection is modelled as a mixture over an underlying set of topics, in which each topic is modelled as a mixture over an underlying set of topic probabilities. These probabilities provide an explicit representation of a document. The algorithm is typically used to extract topics over different documents to classify the documents according to their topics. Here we used the LDA to derive topic keywords that best represent the main idea of each talk across the segmented speech chunks described above. Also, the main idea of a talk is better represented in a phrase, not a single word, for example, “save energy” rather than “save” or “energy”, so we used bi-gram (two words) model in the algorithm to identify topic keywords. For fitting the model, a fixed number of topics should be specified in advance, and we set 4 in the current study.

LDA model represents documents as mixtures of topics with certain probabilities. The assumption of the model posits that documents (speech chunks in the current study) are probability distributions over latent topics (top figure in the third column in Fig. 1b), and topics are probability distributions over words (bi-grams in the current study) in a given corpus (bottom figure in the third column in Fig. 1b). In the model, it is assumed that documents have been produced as follows: First, the number of words in the document is decided; second, a topic mixture is chosen for the document according to a probability distribution over a fixed set of k number of topics which has been set in advance (k=4), for example, 60% of “machine age”, 20% of “industrial revolution”, 10% of “race (with the) machine(s)”, 10% of “productivity time”; third, each word is generated in the document by choosing a topic according to the multinomial distribution in the previous step and using the topic to generate the word based on the topic’s multinomial distribution. This way, the LDA model learns the topic representation of each document and the words associated with each topic. While going through each document, the model randomly assigns each word in the document to one of the k topics. This process results in topic representations of all the documents and word distributions of all topics, albeit initial topics are likely to be inadequate. This step is iterated over every word in every document to improve the model performance.

For every word (bi-gram in the current study) in every document (speech chunk in the current study) and for each topic:

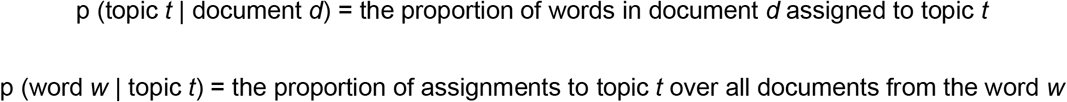

Then the model reassigns *w* a new topic where topic *t* with probability:

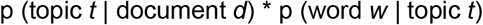

that represents the probability that topic *t* generated word *w*.

Following the process above, documents assigned to topic *t* and words with a high probability for topic *t* are obtained as outputs (document-topic matrix). Here, the most common words (bi-grams in the current study) with the highest probability for topic *t* can be obtained, as shown in Fig. 1c. Also, vectors of topic probabilities for each document (speech chunk in the current study) were depicted using a stacked bar chart (topic mixture), as shown in Fig. 1d.

#### T-SNE representation of LDA result

In addition to a figure for topic mixture showing the probability distribution of each speech chunk over topics, the outputs from the LDA topic model were analyzed using the t-SNE (t-distributed Stochastic Neighbor Embedding) technique (van der Maaten and Hinton, 2008) via scikit-learn (class: sklearn.manifold.TSNE). T-SNE is an unsupervised, nonlinear dimensionality reduction algorithm and is used as a tool for visualizing high-dimensional data by converting pairwise similarities (Euclidean distance as a metric) between data points to joint probabilities by minimizing the Kullback-Leibler divergence (Kullback and Leibler, 1951) between the joint probabilities of the low-dimensional embedding and the original high-dimensional data. Student t-distribution is used for the creation of low-dimensional space instead of a Gaussian distribution for better modelling of distances since t-distribution is heavier-tailed than the Gaussian distribution. Returning values are embeddings of the training data in low-dimensional (2-d) space represented (t-SNE dimensions 1 and 2). The document-topic matrix from LDA topic model analysis was subjected to t-SNE dimensionality reduction model which was then fitted to be visualized in 2-d embedded space. The output from t-SNE technique was visualized using scatterpie chart (Yu, 2020) via ggplot2 (Wickham, 2016) in R (2020), as shown in Fig. 1e. Each data point as a scatterpie chart represents each speech chunk that is clustered into a certain topic *t* according to the topic’s highest probability. In addition, the scatterpie chart shows the probability distribution over topics in a given speech chunk.

### Analysis of neural processing of topic keywords

#### Selection of representative topic keywords

Topic probabilities for each speech chunk were averaged across all speech chunks in a talk in order to identify the representative topic keywords (bi-gram in the current study). For instance, averaging topic probabilities across speech chunks for each topic 1-4 (Fig. 1d) produced topic keywords with the highest topic probability (e.g., topic 3 in this sample talk (see Fig. 1c), “The key to growth? Race with the machines” by Erik Brynjolfsson). Next, the speech chunks were sorted from highest to lowest topic probability as to this topic 3. In order to investigate the neural processing of topic representation, we split these speech chunks into high and low topic probability conditions (Fig. 1f). Corresponding brain data both at sensor and source level were split to high and low topic probability conditions as well, resulting in trial-based epochs, as shown in Fig. 1g.

#### Topic keywords processing of attended vs. unattended talk

The current study aims to identify how the brain processes topic keywords not only in an attended talk, but also in an unattended talk. In the context of natural speech with competing speakers, highly semantic words and sentences in unattended talks can interfere with focused attention to attended talks due to intermittent silences and pauses and semantically less important contents in attended talks, which enables shifting attention to unattended talks. In order to identify the effect of highly semantic speech chunks in unattended talks on the attended speech processing, we performed the same analysis described above for unattended talks as well.

### Global brain activity reflects topic probability in speech chunks

#### Mean-field power analysis at sensor level

Speech chunks with high and low topic probability in both attended and unattended talks were first yielded to mean-field power analysis at the sensor level for a sanity check. The averaged power across left auditory sensors (inset) was depicted in Fig. 2, and effects are shown for each condition (high and low topic probability) for attended and unattended, as well as combined effects for topic probability (regardless of attention) and attention (regardless of topic probability).

**Figure 2.**
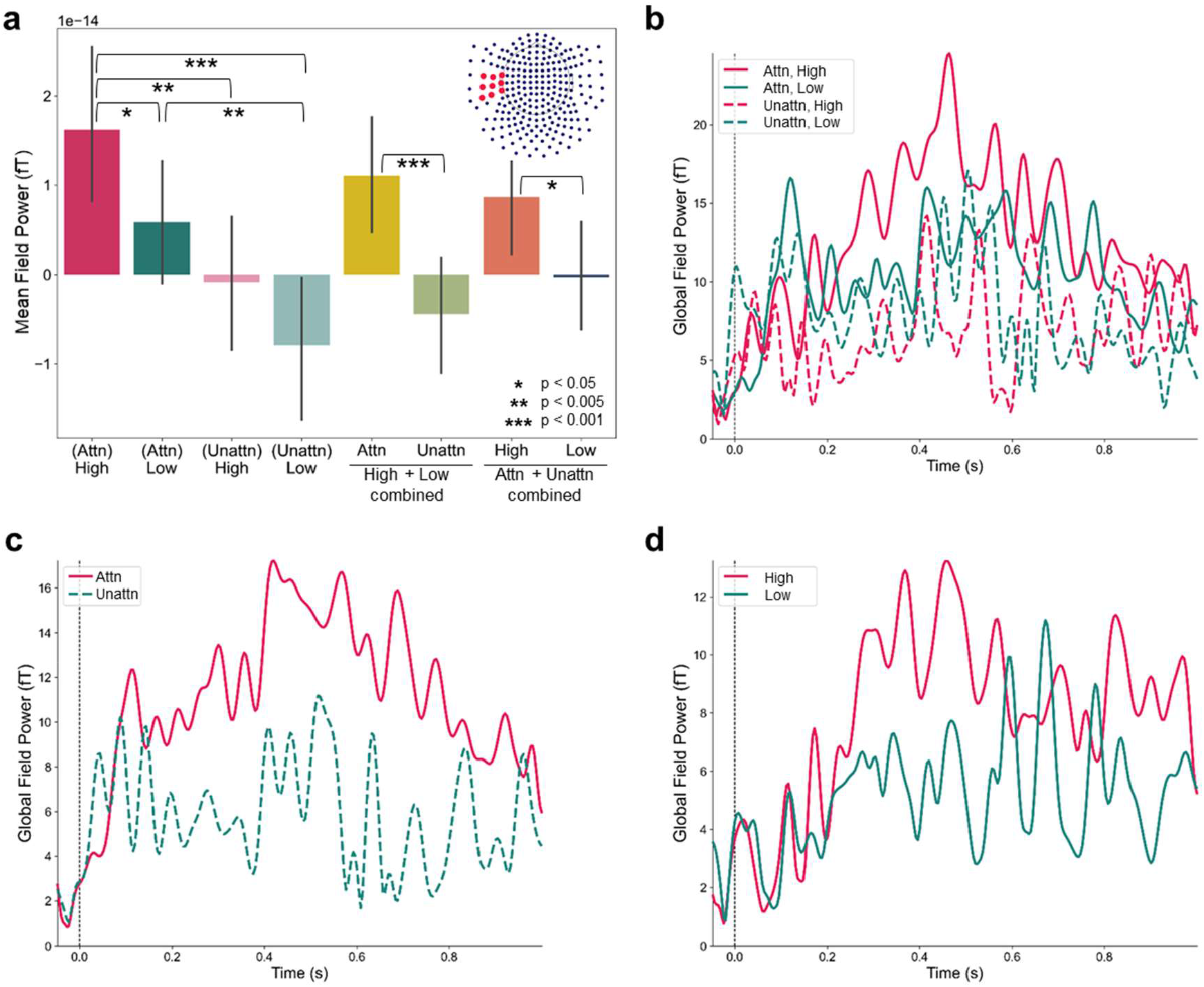
Mean-field power analysis at sensor level. **a**, Mean-field power from 0.25 s to 0.5 s time-locked to the onset of speech chunks over left auditory sensors (inset) were averaged and compared between all combinations of topic probability (high, low) and attention (attended, unattended) as follows via paired t-test: attended, high vs. attended, low: t_43_ = 2.21, p = 0.03; attended, high vs. unattended, high: t_43_ = 3.21, p = 0.002; attended, high vs. unattended, low: t_43_ = 4.35, p < 0.0001; attended, low vs. unattended, high: t_43_ = 1.51, p = 0.14; attended, low vs. unattended, low: t_43_ = 2.91, p = 0.005; unattended, high vs. unattended, low: t_43_ = 1.44, p = 0.16; attended vs. unattended talk pooling across high and low topic probability: t_43_ = 4.17, p = 0.0001; high vs. low pooling across attended and unattended: t_43_ = 2.62, p = 0.01). **b-d**, Temporally unfolded mean-field power by root mean square averaged over the same left auditory sensors during -0.05 s to 1 s time-locked to the speech chunk onset are displayed for all individual conditions (**b**) and combined effects for topic probability (**c**) and attention (**d**).

First, we averaged mean-field power from 0.25 s to 0.5 s time-locked to the onset of speech chunks over left auditory sensors and then statistically compared all combinations of topic probability (high, low) and attention (attended, unattended) (Fig. 2a, paired *t*-test; attended, high vs. attended, low: t_43_ = 2.21, p = 0.03; attended, high vs. unattended, high: t_43_ = 3.21, p = 0.002; attended, high vs. unattended, low: t_43_ = 4.35, p < 0.0001; attended, low vs. unattended, high: t_43_ = 1.51, p = 0.14; attended, low vs. unattended, low: t_43_ = 2.91, p = 0.005; unattended, high vs. unattended, low: t_43_ = 1.44, p = 0.16). In addition, speech chunks from the attended and unattended talk comprising high and low topic probability, i.e., regardless of topic probability, were compared (attended vs. unattended: t_43_ = 4.17, p = 0.0001). Speech chunks from high and low topic probability comprising attended and unattended talk, i.e., regardless of attention, were compared as well (high vs. low topic probability: t_43_ = 2.62, p = 0.01). To elaborate this, we show mean-field power by root mean square in Fig. 2b-d averaged over the same left auditory sensors during -0.05 s to 1 s for each condition (b) and combined effects for topic probability (c) and attention (d).

Speech chunks with high probability from attended talks show the strongest field power. Across both levels of topic probability, attended talk shows stronger field power than unattended talk (Fig. 2c). Across both levels of attention, speech chunks with high topic probability show stronger mean field power than speech chunks with low topic probability (Fig. 2d). The peak around 0.4 s in Fig. 2b-d is likely to reflect meaning processing in the brain (Kutas and Federmeier, 2011).

### Receptive field model estimation for neural topic keywords processing

We next implemented a receptive field model, also known as a multivariate temporal response function (mTRF) (Crosse et al., 2016), which can be interpreted as a linear filter in the brain processing a stimulus feature (speech envelope S(t)) mapped onto the continuous neural responses (MEG response over time, r(t)). The approach has been used in recent studies to study the mapping between brain responses and naturalistic speech feature representations (Broderick et al., 2018; Teng et al., 2021; Haider et al., 2022). The schematic flowchart for our analysis is shown in Fig. 2a, which was adapted from Fig. 1 in Crosse et al. (2016).

The stimulus feature-neural response mapping can be modelled bidirectionally (forward and backward modelling). In the encoding (forward: stimulus to brain) model, the model function mathematically describes how the speech amplitude envelope is encoded in MEG responses which can be interpreted as underlying neural generators (Crosse et al., 2016). To derive the encoding model estimation, a time-lagged (−0.2 s to 0.8 s in steps of 8 ms) speech envelope from each epoch in each condition (high, low topic probability in each attended and unattended talk) was used as an input feature to predict corresponding neural responses (248 sensor MEG response).

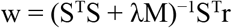

The fitting of the encoding model implemented the ridge regression (also known as Tikhonov regularization), where the loss function is the linear least-squares function and regularization (ridge parameter λ) is given by the L2-norm. The estimate is improved by using quadratic penalization (M) of the difference between two neighbouring terms of w (Lalor et al., 2006). The K-fold (k=3) cross-validation was used to split data into train and test sets (5:5), and fitting the model was iterated through the subset. This computation was performed via MNE-Python (Gramfort et al., 2013) class mne.decoding.ReceptiveField and separately for each condition (attended, high; attended, low; unattended, high; unattended, low). Model coefficients maps (weight vectors from the model estimation; TRF w, number of cross-validation x number of sensors x number of time delays: 3 × 248 × 126) were obtained and averaged over cross-validation splits. The prediction score (derived as the correlation coefficient, *r*) between predicted and original neural responses was also computed (number of cross-validation × number of sensors) and averaged across cross-validation splits.

We performed the decoding model (backward: brain to stimulus; speech reconstruction) estimation as well (see for more details in Supplementary Figure 2) and investigated model coefficients map and prediction scores between conditions of combinations of attention (attended, unattended) and topic probability (high, low). The prediction score (derived as the correlation coefficient, *r*) between the reconstructed and original speech envelope was computed and averaged across cross-validation splits, as shown in Fig. 3b. One representative result for the reconstructed speech envelope is shown on the left bottom in Fig 3a.

**Figure 3.**
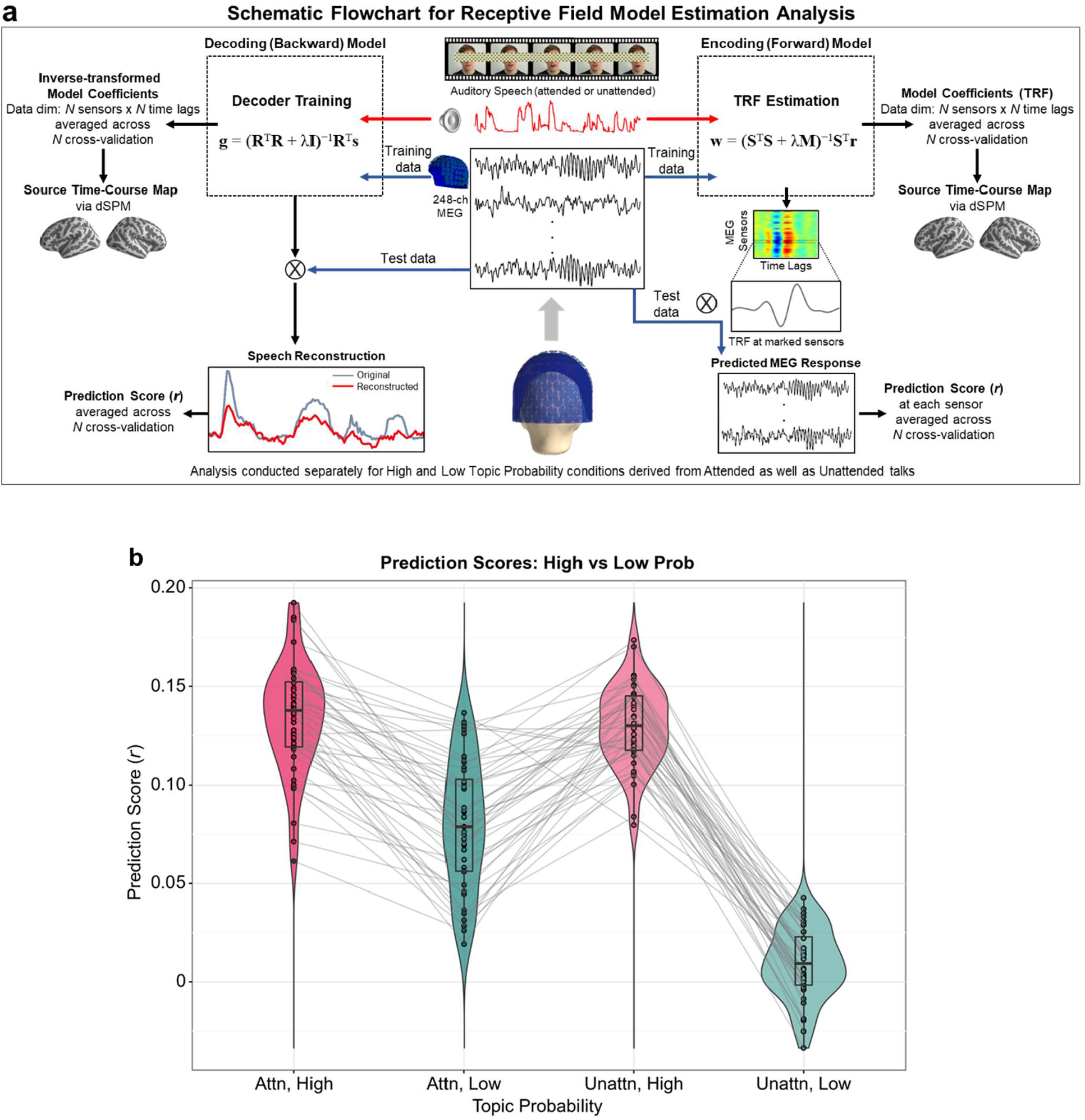
Receptive field model estimation analysis for topic probability and attention effects. a, Schematic flowchart for the model estimation. The stimulus feature-neural response mapping was modelled bidirectionally using encoding and decoding models (also known as forward and backward models). Encoding and decoding model analyses were performed separately for high and low topic probability conditions as well as for attended and unattended talks. Figure adapted from Crosse et al. (2016). **b, Decoding model performance**. Model prediction accuracy to reconstruct speech envelope for each condition was obtained by correlation coefficient score (*r*-value) between original and reconstructed speech, which were then compared between high and low topic probability conditions separately for each attended and unattended talk. The prediction accuracy was significantly stronger for speech with high than low topic probability in both attended (between 1^st^ and 2^nd^ columns: attended, high vs. attended, low; t_43_ = 11.43, p = 1.27e-14) and unattended talk (between 3rd and 4th columns: unattended, high vs. unattended, low; t_43_ = 29.98, p = 1.88e-30) with a greater difference for unattended than attended talk. Interestingly, the prediction accuracy was stronger for the high topic probability condition in unattended talk than the low topic probability condition in attended talk (between 2nd and 3rd columns: unattended, high vs. attended, low; t_43_ = 9.62, p = 2.74e-12). However, no significant difference was observed between high topic probability conditions (between 1st and 3rd columns: attended, high vs. unattended, high; t_43_ = 0.80, p = 0.43). All statistical comparisons were performed via two-tailed paired t-test. Dots and lines represent individual results.

For the investigation of model coefficients map at the source level, dynamic statistical parametric mapping (dSPM) (Dale et al., 2000; Gramfort et al., 2014) was applied as an inverse solution. The computation was performed separately for each condition, and the statistical significance maps between the conditions were derived. The results from the encoding model map, as summary clusters, are shown in Fig. 4, and the results for significant temporal clusters from the encoding model can be found in Supplementary Figure 1. Decoding model weights are not interpretable in a neurophysiological sense due to potential spurious observations; however, inverse-transformed decoder weights, i.e., transforming the backward model into the forward model via pseudo-inverse, can provide physiologically interpretable information (Haufe et al., 2014). For this work, please see Supplementary Figure 2.

**Figure 4.**
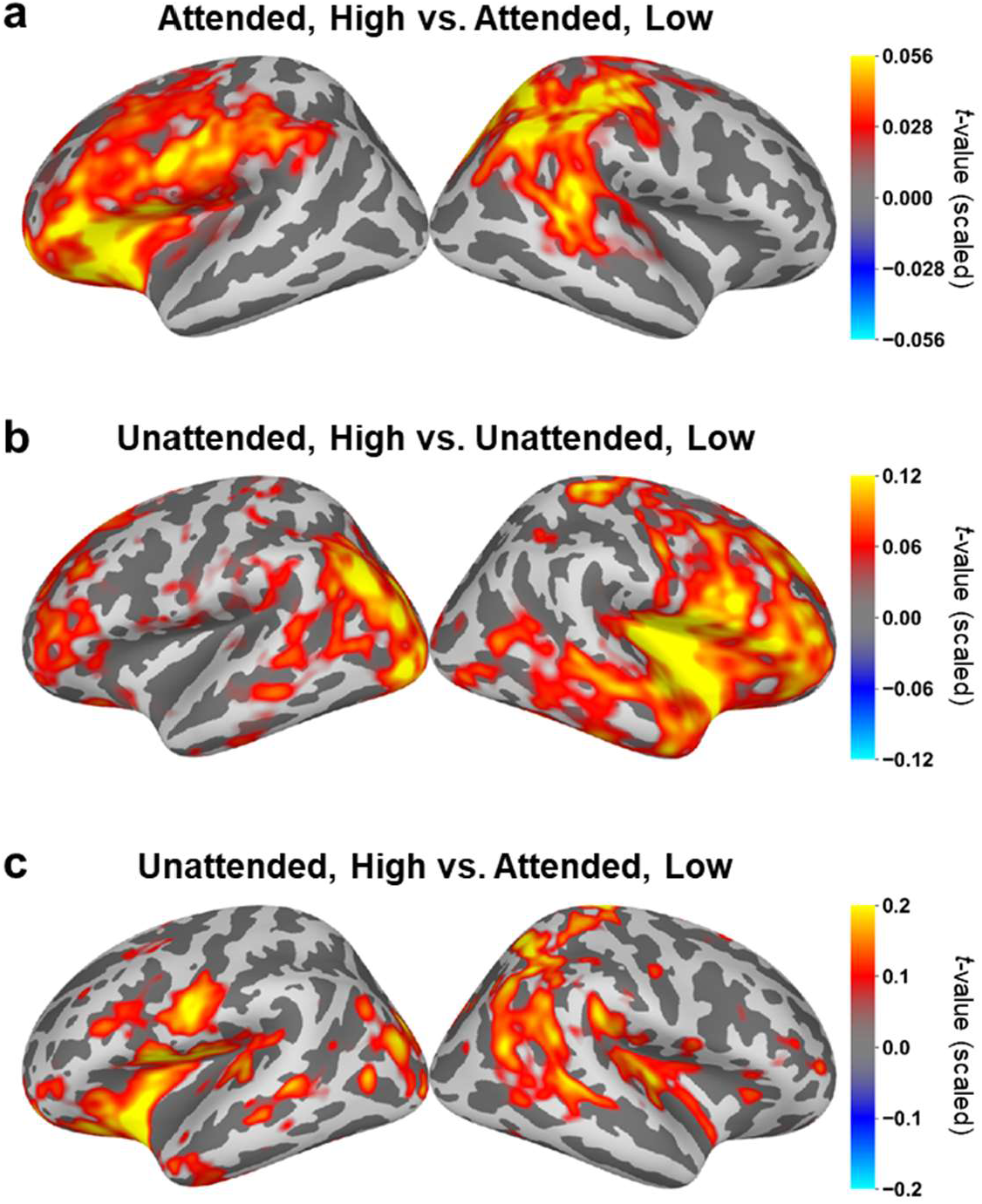
Encoding model weights mapped onto source space. Encoding model coefficients were mapped onto source space via dSPM method and statistically compared between high and low topic probability conditions using cluster-level spatio-temporal permutation test (p < 0.05; two-tailed; 1024 permutations). Summary clusters (averaged across all significant temporal clusters) are shown. **a**, attended, high vs. attended, low. **b**, unattended, high vs. unattended low. **c**, unattended, high vs. attended, low. T-values in significant clusters are scaled corresponding to the duration spanned by the cluster (for more details, see Statistical test in Materials and Methods).

### Stronger speech reconstruction accuracy for speech chunks with high than low topic probability

The decoder model can provide a complementary view on the interpretation of the stimulus feature-neural response relationship as follows. First, the decoder can be used to reconstruct stimulus features from the neural responses. In the current study, a trained decoder was used to reconstruct the speech envelope. The prediction accuracy to reconstruct speech, defined as correlation coefficient score (*r*) between original and reconstructed speech envelope, was significantly stronger for speech with high than low topic probability in both attended (attended, high vs. attended, low: paired t-test; t_43_ = 11.43, p = 1.27e-14; between 1^st^ and 2^nd^ columns in Fig. 3b) and unattended talk (unattended, high vs. unattended, low: paired t-test; t_43_ = 29.98, p = 1.88e-30; between 3^rd^ and 4^th^ columns in Fig. 3b). The difference between high and low topic probability speech is greater for speech chunks derived from unattended talk when compared to attended talk. Furthermore, speech reconstruction performance is stronger for high topic probability condition in unattended talk than low topic probability condition in attended talk (unattended, high vs. attended, low: paired t-test; t_43_ = 9.62, p = 2.74e-12; between 2^nd^ and 3^rd^ columns in Fig. 3b). This implies that topic keywords in unattended talks are still captured and are processed to the same degree as those in attended talks. No significant difference was observed between high topic probability conditions (attended, high vs. unattended, high: paired t-test; t_43_ = 0.80, p = 0.43; between 1^st^ and 3^rd^ columns in Fig. 3b). All other paired t-test results between conditions are as follows: attended, high vs. unattended, low: t_43_ = 22.61, p = 1.71e-25; between 1^st^ and 4^th^ columns and attended, low vs. unattended, low: t_43_ = 11.79, p = 4.62e-15; between 2^nd^ and 4^th^ columns in Fig. 3b).

Next, we investigated the spatial representations of topic keywords processing between high and low topic probability condition pairs that showed significant prediction performance above (attended, high vs. attended, low; unattended, high vs. unattended, low; unattended, high vs. attended, low). The encoding model coefficients for each condition were mapped on the brain surface using the dynamic statistical parametric mapping (dSPM) source localization method. The statistical difference was derived via two-tailed cluster-level spatio-temporal permutation t-test (p < 0.05; 1024 permutations) for 0 to 0.5 s with respect to the speech chunk onset in steps of 0.008 s in MNE-Python (function: mne.stats.spatio_temporal_cluster_1samp_test). Here we show summary clusters, i.e., all significant clusters pooled across the temporal clusters. For temporally unfolded clusters, including before speech onset (−0.2 to 0 s), please see Supplementary Figure 1.

For attended speech (Fig. 4a), the difference between high and low topic probability was observed in the left inferior frontal (BA 44, 45), somatosensory areas, as well as primary motor (BA 4), premotor (BA 6) cortices and right supramarginal, angular and posterior superior temporal gyri. For unattended speech (Fig. 4b), the difference between high and low topic probability was observed in the right inferior frontal areas, insula and temporal areas and left frontal, posterior temporal and visual cortex. We further performed the same statistical test between unattended high vs. attended low topic probability conditions where significant speech reconstruction performance was observed. The result (Fig. 4c) showed similar brain regions as to attended high vs. attended low (Fig. 4a), though to a lesser extent, suggesting speech chunks with high topic probability in unattended talks are processed to a similar degree as those in attended talks.

### Semantically salient speech in unattended talk causally mediates attended speech comprehension

As shown above, the statistical difference between high and low topic probability was greater for unattended talks than for attended talks. Furthermore, the difference was evident for unattended but with high topic probability speech chunks when compared to attended but with low topic probability condition. These findings suggest that distracting speech chunks are still processed when they contain semantic gist. This has led us to hypothesize that attention to semantically salient unattended speech negatively mediates (i.e., suppresses) attended speech comprehension. We employed the mediation analysis (Baron and Kenny, 1986) to inspect the relationship between unattended speech and attended speech comprehension. Encoding model coefficients for 0.5 s after the onset of speech chunks were averaged within each parcellation of the PALS-B12-Brodmann atlas (Van Essen, 2005) within each individual subject. The PALS-B12-Brodmann atlas parcellation has 82 cortical structures with 41 structures for each hemisphere. In the mediation model, encoding model coefficients of attended speech chunks with high topic probability, encoding model coefficients of unattended speech chunks with high topic probability, and speech comprehension accuracy for attended speech were used as predictor (*X*, independent), mediator (*M*), and target (*Y*, dependent) variables, respectively.

We found significant negative indirect effects for 4 out of 82 cortical regions: BA41 (primary auditory cortex), BA4 (primary motor cortex), BA6 (premotor cortex) in the left hemisphere and BA9 (dorsolateral prefrontal cortex) in the right hemisphere (Left BA41: path ab β = -2.03, p = 0.03, 95% CI = -5.53, -0.29; Left BA4: path ab β = -3.82, p = 0.02, 95% CI = -11.12, -0.71; Left BA6: path ab β = -3.51, p = 0.009, 95% CI = -10.84, -0.93; Right BA9: path ab; β = -6.43, p = 0.03, 95% CI = -13.44, -0.95; 5000 bootstrap iterations performed; Fig. 5a). There were no significant direct effects. As such, the findings support the complete (full) mediation effects (also referred to as causal mediation effect), indicating the increased activities in these brain regions to unattended (*to-be-ignored*) speech suppress attention to attended (*to-be-attended*) speech leading to poor speech comprehension performance. The encoding model coefficients in these regions for all conditions including low conditions are shown in Supplementary Figure 3.

**Figure 5.**
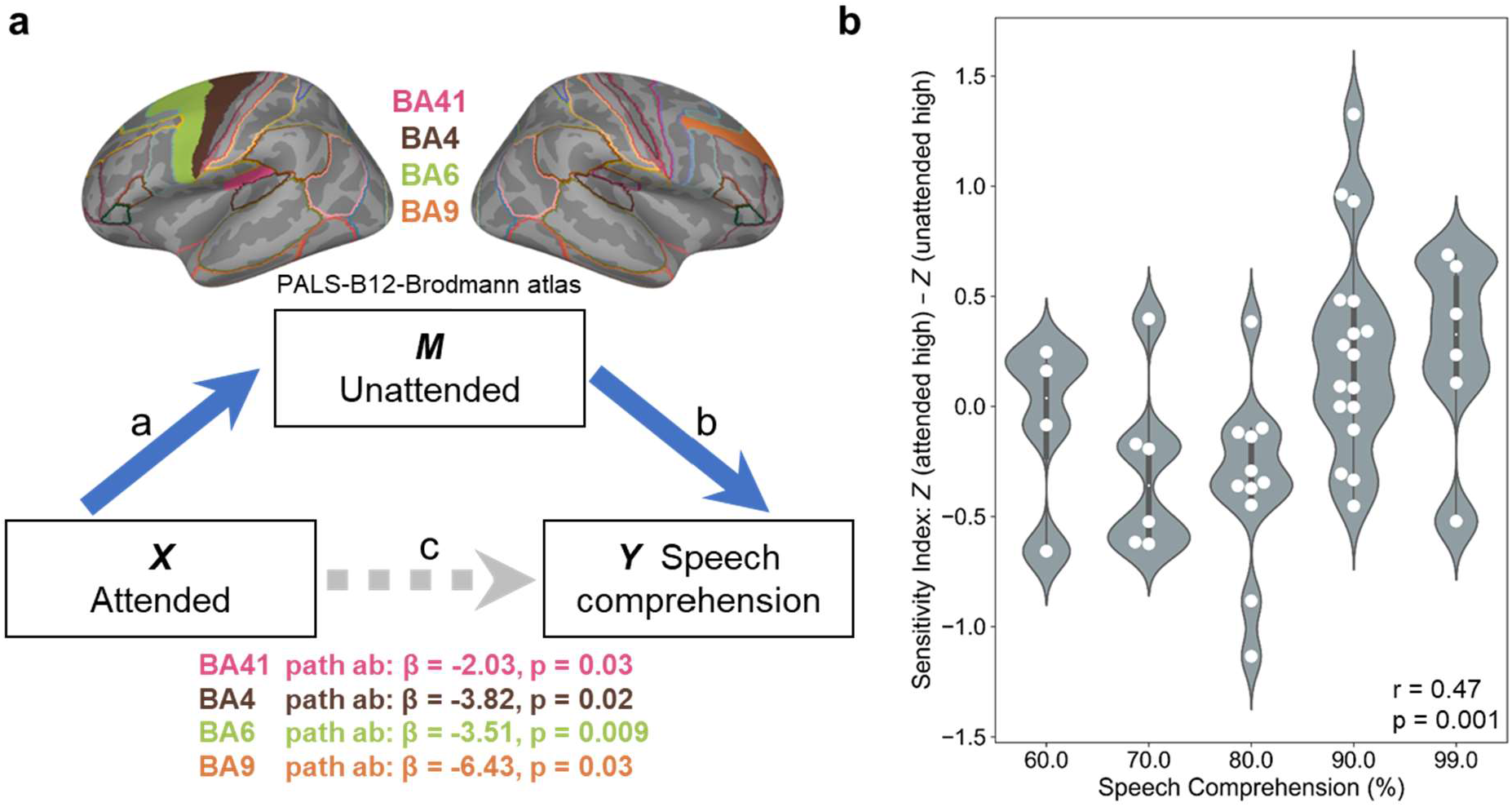
Causal relationship between attended and unattended speech on speech comprehension. a, Salient unattended speech negatively mediates attended speech comprehension. Mediation analysis was performed to test the hypothesis that attention to semantically salient unattended speech negatively mediates (i.e., suppresses) attended speech comprehension. In the mediation model, encoding model coefficients of attended speech chunks with high topic probability, encoding model coefficients of unattended speech chunks with high topic probability, and speech comprehension accuracy for attended speech were used as predictor (*X*, independent), mediator (*M*), and target (*Y*, dependent) variables, respectively. The encoding model coefficients during 0 - 0.5 s with respect to the speech chunk onset were averaged within each region in PALS-B12-Brodmann atlas in MNE-Python within each individual. Significant negative indirect effects were identified for left BA41 (path ab β = -2.03, p = 0.03, 95% CI = -5.53, -0.29), left BA4 (path ab β = -3.82, p = 0.02, 95% CI = -11.12, -0.71), left BA6 (path ab β = -3.51, p = 0.009, 95% CI = -10.84, -0.93) and right BA9: path ab; β = -6.43, p = 0.03, 95% CI = -13.44, -0.95) with 5000 bootstrap iterations. **b, Increased sensitivity to attended speech in the regions enhances speech comprehension**. Sensitivity index, analogous to d-prime, defined by the difference between *Z*-transformed model coefficients of attended speech with high topic probability and unattended speech with high topic probability (Z (attended high) – Z (unattended high)) was created for each of 4 regions and averaged. The sensitivity index was significantly correlated with speech comprehension accuracy across participants (Spearman rank correlation: r = 0.47, p = 0.001), supporting the hypothesis that participants with increased sensitivity to semantically salient attended speech in these regions show better speech comprehension.

To substantiate this modulatory effect, we tested if increased sensitivity to attended speech in these regions is associated with better speech comprehension. For this, we made a sensitivity index using the difference between *Z*-transformed coefficients of attended speech with high topic probability and unattended speech with high topic probability (*Z* (attended, high) – *Z* (unattended, high)), analogous to the d-prime (i.e., sensitivity index, also known as discriminability or detectability). Then, the sensitivity index values were averaged across the four brain regions and correlated with speech comprehension accuracy using Spearman rank correlation over participants (r = 0.47, p = 0.001 in Fig. 5b). The result supports that participants with increased sensitivity to attended (*to-be-attended*) talks with high topic probability in these brain regions indeed show better speech comprehension.

## Materials and Methods

### Participants

Forty-six native English speakers participated in the study. All participants reported normal hearing (confirmed by two hearing tests using research applications on an iPad: uHear (Unitron Hearing Limited) and Hearing-Check (RNID)) as well as normal or corrected-to-normal vision. They all had no history of neurological, developmental, or psychological disorders and were all right-handed, confirmed by the Edinburgh Handedness Inventory (Oldfield, 1971). Data from 44 subjects were analyzed (26 females; age range: 18–30 y; mean age: 20.54 ± 2.58 y) after two subjects were excluded since one subject fell asleep and one had excessive signal noise). Other analyses of these data were presented in previous reports (Park et al., 2016; Park et al., 2018b). All subjects provided informed written consent before the experiment and received monetary compensation for their participation. The study was approved by the local ethics committee (CSE01321; College of Science and Engineering, University of Glasgow) and undertaken in accordance with the ethical guidelines in the Declaration of Helsinki.

### Stimuli and Experiment

#### Natural speech materials

Each auditory speech presented to the participants during MEG recordings was about a certain coherent topic, and the materials we presented to the participants were originally taken from TED talks (www.ted.com/talks/) and modified to be appropriate for our own filming (e.g., “The key to growth? Race with the machines” by Erik Brynjolfsson. Transcription for each talk was downloaded and edited to be appropriate for our own filming by editing words such as referring to visual materials, the gender of the speaker etc. The talks address a specific topic belonging to informative, persuasive, inspiring categories on the website. Please note that they no longer provide these categories explicitly. We additionally validated the speech materials in a separate behavioural study (33 participants with 19 females; aged 18–31 years; mean age: 22.27 ± 2.64 years), where participants rated the talks in terms of arousal, familiarity, valence, complexity, significance (informativeness), agreement (persuasiveness), concreteness, self-relatedness, and level of understanding using Likert scale (Likert, 1932) 1 to 5 (for an example of concreteness, 1: very abstract, 2: abstract, 3: neither abstract nor concrete, 4: concrete, 5: very concrete). Talks with excessive mean scores (below 1 and over 4) were excluded, and eight out of eleven videos were selected for the experiment, and selected talks were used in different experimental conditions, which were also used in our previous study (Park et al., 2016). High-quality audiovisual video clips were filmed by a professional filming company while a professional male speaker was talking continuously. The duration of talks is 7 to 9 minutes with a sampling rate of 48 kHz for audio and 25 frames per second (fps) for video in 1,920 × 1,080 pixels. Filmed videos were edited for experimental manipulations (i.e., recombinations of auditory and visual speech stimuli) using Final Pro Cut × (Apple, Cupertino, CA).

#### Experimental condition

We employed four experimental conditions as described in our previous study (Park et al., 2016). In the current study, we focused on the “AV congruent” condition where two different talks are delivered to the left and right ear with one auditory speech matching the visual lip movement, and the speech presented to the other ear serves as a distractor. Participants were instructed to pay attention to the talk that matches visual lip movement. The side of attention was counterbalanced, resulting in half of the participants (N = 22) paying attention to the left-ear talk, whereas the other half (N = 22) paying attention to the right-ear talk. Participants were instructed to fixate on the visual information, i.e., the speaker’s lip movement and subjects’ eye movements were monitored using an eye tracker (EyeLink 1000, SR Research). To assist this, a fixation cross color-coded either yellow or blue was overlaid on the speaker’s lip, and the color of the fixation cross indicated the side of attention. During the experiment, the audiovisual stimuli were presented via Psychtoolbox (Brainard, 1997) in MATLAB (MATLAB, R2019b). Auditory stimuli were delivered at a 48 kHz sampling rate via a sound pressure transducer through 2 five-meter-long plastic tubes terminating in plastic insert earpieces, and visual stimuli were presented with a resolution of 1,280 × 720 pixels at 25 fps (mp4 format).

#### Behavioural performance

In order to assess the level of comprehension of the talk, a questionnaire for speech comprehension was administered after the talk. The questionnaire consists of 10 questions about the attended talk, such as “What is the speaker’s job?” and “What would be the best title of this talk?”. The questionnaire itself was validated in a separate behavioural study (16 subjects; 13 females; aged 18–23 y; mean age: 19.88 ± 1.71 y) in terms of accuracy (the same level of difficulty), response time, and the length (word count). The comprehension scores for left- and right-ear attention group did not differ (two-sample t-test for “attended to left” vs. “attended to right” group, t_42_ = -0.13, p = 0.90). In this study, we pooled across both groups in all data analyses so that attentional effects for a particular side (e.g., left or right) are expected to cancel out.

### MEG and MRI (T1) data processing

#### Data acquisition

Neuromagnetic signals were measured using a 248 magnetometers whole-head MEG system (MAGNES 3600 WH, 4-D Neuroimaging) in a magnetically shielded room with a sampling rate of 1,017 Hz. The signals were resampled to 250 Hz and denoised with information from the reference sensors using the denoise_pca function in the FieldTrip toolbox (Oostenveld et al., 2011). Bad MEG sensors were excluded by visual inspection, and electrooculographic (EOG) and electrocardiographic (ECG) artefacts were rejected using independent component analysis (ICA). In order to map MEG data onto cortical source space, structural T1-weighted MRIs of each participant were acquired at 3 T Siemens Trio Tim scanner (Siemens, Erlangen, Germany) with the following parameters: 1.0 × 1.0 × 1.0 mm^3^ voxels; 192 sagittal slices; field of view (FOV): 256 × 256 matrix.

#### Coregistration between MRI and MEG data

Structural T1 MR images recorded from each participant were coregistered to the MEG coordinate system via a semiautomatic procedure. Anatomical fiduciary landmarks such as nasion and bilateral preauricular points were identified before the MEG recording and also manually identified in the individual’s MR images. Based on these landmarks, both MEG and MRI coordinate systems were initially aligned, followed by numerical optimization achieved by using the ICP algorithm (Besl and McKay, 1992).

#### Source localization

Reported results in the current study were analyzed in both Fieldtrip (Oostenveld et al., 2011) and MNE-Python (Gramfort et al., 2014), so workflow for inverse solution followed the standard procedure in each software. In Fieldtrip, a head model was created for each individual from their structural MRI using normalization and segmentation routines in FieldTrip and SPM8. For the calculation of the leadfield, we used a single-shell volume conductor model (Nolte, 2003) employing an 8 mm grid defined on the template brain provided by MNI (Montreal Neurological Institute). The template grid was linearly transformed into individual headspace for spatial normalization. In MNE-Python, construction of the forward model solution and MRI segmentation are performed in the FreeSurfer package (Dale et al., 1999). The Boundary Element Model (BEM) of individual MRI was created using the FreeSurfer watershed algorithm, and surface-based source space with a 4.9 mm source spacing resolution was computed. Then individual source map was morphed to the template MRI (fsaverage) to compare output activities across subjects in common source space.

### Audiovisual speech signal processing

#### Auditory speech signal

The amplitude envelope of sound signals was computed following the approach introduced in Chandrasekaran et al. (2009). We first constructed eight frequency bands in the range of 100–10,000 Hz to be equidistant on the cochlear map (Smith et al., 2002). Then the auditory sound speech signals were band-pass filtered in these bands using a fourth-order forward and reverse Butterworth filter followed by Hilbert transform to obtain amplitude envelopes for each band of the signal. These signals were then averaged across bands resulting in a wideband amplitude envelope. Signals were downsampled to 250 Hz for further analysis to match the sampling rate of preprocessed MEG data.

#### Visual speech signal

A lip movement signal was computed using an in-house MATLAB script as in our previous report (Park et al., 2016), which demonstrated oscillatory brain activities entrained by visual speech. We first extracted the outline lip contour of the speaker for each frame of the video stimuli. From the lip contour outline, we computed the frame-by-frame lip area (area within lip contour). This signal was resampled at 250 Hz to match the sampling rate of the preprocessed MEG signal and auditory sound envelope signal.

### Mediation analysis

We used mediation analysis (Baron and Kenny, 1986) to test our hypothesis that attention to semantically salient unattended speech negatively mediates (i.e., suppresses) attended speech comprehension using mediation_analysis module in Pingouin package (Vallat, 2018). Here we used predefined cortical parcellation, the PALS-B12-Brodmann atlas (Van Essen, 2005). The parcellation provides 82 cortical surface structures (41 in each hemisphere). Encoding model coefficients (TRF) is averaged across 0 s to 0.5 s after the onset of speech chunks within all the areas in the PALS-B12-Brodmann parcellation for each subject. In the mediation model, these values of attended speech chunks with high topic probability and unattended speech chunks with high topic probability were used as predictor (X, independent) and mediator (M) variables, respectively. For the dependent variable (Y), speech comprehension accuracy for attended speech was used. Five thousand bootstrap iterations were performed for confidence intervals and p-values estimation. To confirm the mediation effects, we performed correlational analysis between the sensitivity index, defined by the difference between z transformed attended high (analogous to hit) and unattended high (analogous to false alarm) conditions, and speech comprehension accuracy.

### Statistical test

All the analyses described in the Methods section were performed individually and then yielded to group-level statistical tests on the data of all 44 participants.

For receptive field model estimation (mTRF), the non-parametric cluster-level paired permutation test based on a t-statistic (Maris and Oostenveld, 2007) was performed at the spatio-temporal level (function: mne.stats.spatio_temporal_cluster_1samp_test) between high vs. low topic probability conditions after morphing into common cortical space (fsaverage) in MNE-Python (Gramfort et al., 2013). The function detects significant clusters at both spatial and temporal regions. A spatial adjacency matrix was used for clustering in source space, and 1024 permutations were computed. T-values in significant clusters are scaled corresponding to the duration spanned by the cluster (shown as “t-value (scaled)” in figures) in which the scaling factors are the significance of the t-value and the unit of time step between samples (e.g., 0.008 s) in the data. In more detail, the significance of t-values at a given time point (1 if significant, 0 if not) is scaled by the unit of the duration (e.g., significance (1 or 0) × 0.008 s). These values are summed up across significant time points. Each temporal cluster from the encoding model statistics is shown in Supplementary Figure 1.

## Discussion

In the current study, we used computational topic modelling to investigate how the brain processes high-level semantics in naturalistic speech. Topic modelling techniques, such as Latent Dirichlet Allocation (LDA), have broad application not only to text materials but to other domains, for example, content-based images (Blei et al., 2003). The LDA technique provides scalable and quantifiable measures in identifying topics using a generative probabilistic approach. To date, studies of neural mechanisms underlying semantic processing during natural speech perception have largely relied on the word level violations or predictions (Kutas and Federmeier, 2011; Wang et al., 2018; Broderick et al., 2019; Koskinen et al., 2020). Investigating the neural representation of latent word meanings, such as arising from idiomaticity, or semantic gist (i.e., topic keywords) across supporting contextual information in a connected speech, requires moving beyond inspecting lexico-semantic level processing; however, it has been challenged due to the lack of appropriate quantifiable approaches. Here we harness the state-of-the-art machine learning-based natural language processing (NLP) algorithm to delineate the neural signatures in topic keywords processing. To our knowledge, this is the first study that investigates neural signatures of high-level semantics processing beyond lexico-semantics in continuous speech.

By applying the topic model to speech chunks that were driven in a perceptually relevant manner by acoustic silences in natural speaking, we were able to extract representative topic keywords of a certain story. Subsequently, speech chunks were split into two statistical conditions according to the probability of the main topic keywords. Corresponding epochs of brain activities were grouped to the conditions. Here we focused our analysis on the topic keywords processing in the context of the multi-speaker environment where the attention to a particular talk is manipulated. Brain responses and stimulus feature (speech envelope) were used to fit the encoding and decoding model to investigate the neural map of semantic gist throughout the speech.

Our finding showing stronger brain activity for attended speech than unattended speech (Fig. 2) is consistent with other findings that showed brain responses to unattended speech is attenuated (Kong et al., 2014). However, strikingly, the difference in reconstruction of speech envelope from neural activities between high and low topic probability conditions in unattended speech is greater when compared to the same difference in attended speech. The finding is even striking for the significant difference between unattended high vs. attended low condition. This result supports the notion that unattended (i.e., task-irrelevant) speech is still processed in the brain due to the failure of selective attention to fully suppress distracting sensory input (Har-Shai Yahav and Zion Golumbic, 2021). Despite decades of debate (Bronkhorst, 2015), a recent study by Har-Shai Yahav and Zion Golumbic (2021) has shown that the phrasal structure of structured task-irrelevant stimuli was represented in the neural responses and competed with task-relevant (attended) speech. Our finding extends this significantly by showing that this is even the case for higher-level semantic processing, indicating that the human brain can capture the semantic gist of the task-irrelevant speech in a multi-speaker context.

We report source mapping using encoding model coefficients via the dSPM source reconstruction method. For attended speech (Fig. 4a), the patterns of the difference between high vs. low topic probability conditions are mapped onto the left inferior frontal, dorsolateral prefrontal and extensive temporal areas corresponding to the language network. For the unattended speech (Fig. 4b), the difference was mapped onto the right inferior frontal/insular areas and the left superior/dorsolateral prefrontal areas. The prominent involvement of the right inferior frontal and insular cortex might suggest lower-level processing of incoming but task-irrelevant speech. While the left hemisphere is actively engaging in semantic processing of goal-directed attended speech, the right hemisphere processes perceptual features such as pitch contours of unattended speech consistent with the notion of the division of labour between hemispheres (Flinker et al., 2019). In Flinker et al. (2019), the authors suggest differential hemispheric contributions in auditory processing in which right lateralization for spectral modulation, such as gender identification, while left lateralization for intelligibility task (Fig. 5 in Flinker et al. (2019)). As such, we suggest that this finding might reflect processing towards a more perceptual level, e.g., pitch contours, with a potential shift of attentional state leading to shallow level semantic processing given the complex nature of dichotic listening (multi-speaker) environment.

Our mediation analysis allows us to further investigate the causal relationship between attended and unattended speech on speech comprehension. The negative full mediation effect (Fig. 5a) indicates that the failure to suppress salient distractors of unattended speech leads to poor speech comprehension of attended speech. Interestingly this effect was observed in the left BA41 (primary auditory cortex), BA4 (primary motor cortex), BA6 (premotor cortex) and right BA9 (dorsolateral prefrontal cortex), which are sub-regions of the speech processing network. Particularly, intrinsic coupling between the auditory and motor system has been previously reported (Assaneo and Poeppel, 2018). However, the role of motor and premotor cortices in speech perception is controversial (Meister et al., 2007; Hickok, 2010; Skipper et al., 2017), and the contribution to the processing of high-level semantics is not well-known. In our previous results using the same dataset, however, the motor cortex was involved in conveying greater information than the linear summation of individual auditory and visual perceptual information (i.e., synergistic information processing) and supported behavioural performance (Park et al., 2018b). Given additional manipulations for high-level semantic processing and attention in the current study, we interpret the motor and premotor cortex might support top-down modulated active sensing (Morillon et al., 2015; Park et al., 2015) for semantic gist in temporally rapidly changing and dynamically evolving naturalistic speech perception in multi-speaker environment. Shifting between task-relevant and -irrelevant semantic gist is suggested to be modulated by these regions, as shown in the result that subjects with increased sensitivity to semantic gist of goal-directed attended speech exhibit better behavioural performance (Fig 5b).

Further investigation will be required to gain deeper insights into high-level semantic processing by fully characterizing the difference from the neural mechanisms underlying other speech features, including lexico-semantic processing, which is expected to provide a direct computational link between our high-level semantic approach and previous work focused on word similarity using other NLP algorithms such as word2vec (Mikolov et al., 2013) as used in Broderick et al. (2018), or semantic neural representation shown in Huth et al. (2016). Furthermore, future studies should be able to provide evidence of how the brain builds up the semantic core with time as the supporting building blocks of different levels of semantic features are accumulated.

In summary, we provide evidence that temporal and spatial neural signatures for high-level semantic gist (i.e., topic keywords) processing in the context of multi-speaker environment and how the semantic gist in unattended speech affects attended speech comprehension.

## Funding

Wellcome Trust Senior Investigator Award https://wellcome.ac.uk/ (grant number 098433) and DFG project funding (GR 2024/5-1, GR 2024/11-1) to JG. The funder had no role in study design, data collection and analysis, decision to publish, or preparation of the manuscript.

## Conflict of interest statement

The authors declare that the research was conducted in the absence of any commercial or financial relationships that could be construed as a potential conflict of interest.

## Notes

### Competing Interest Statement

The authors have declared no competing interest.

## References

Assaneo MF, Poeppel D (2018) The coupling between auditory and motor cortices is rate-restricted: Evidence for an intrinsic speech-motor rhythm. Science Advances 4.

Baron RM, Kenny DA (1986) The moderator-mediator variable distinction in social psychological research: conceptual, strategic, and statistical considerations. J Pers Soc Psychol 51:1173–1182.

Besl PJ, McKay ND (1992) A method for registration of 3-D shapes. IEEE T Pattern Anal:239–256.

Biau E, Wang D, Park H, Jensen O, Hanslmayr S (2021) Auditory detection is modulated by theta phase of silent lip movements. Current Research in Neurobiology 2.

Bird S, Ewan K, Loper E (2009) Natural Language Processing with Python: O’Reilly Media, Inc. Blei DM Topic modeling. In: http://www.cs.columbia.edu/~blei/topicmodeling.html.

Blei DM (2012) Probabilistic Topic Models. Commun Acm 55:77–84.

Blei DM, Ng AY, Jordan MI (2003) Latent Dirichlet allocation. J Mach Learn Res 3:993–1022.

Boersma P, Weenink D (2018) Praat: doing phonetics by computer [Computer program]. Version 6.0.37:retrieved 14 March 2018 from http://www.praat.org/.

Brainard DH (1997) The Psychophysics Toolbox. Spatial vision 10:433–436.

Broderick MP, Anderson AJ, Lalor EC (2019) Semantic Context Enhances the Early Auditory Encoding of Natural Speech. J Neurosci 39:7564–7575.

Broderick MP, Anderson AJ, Di Liberto GM, Crosse MJ, Lalor EC (2018) Electrophysiological Correlates of Semantic Dissimilarity Reflect the Comprehension of Natural, Narrative Speech. Curr Biol 28:803–809 e803.

Bronkhorst AW (2015) The cocktail-party problem revisited: early processing and selection of multi-talker speech. Atten Percept Psychophys 77:1465–1487.

Chandrasekaran C, Trubanova A, Stillittano S, Caplier A, Ghazanfar AA (2009) The natural statistics of audiovisual speech. PLoS computational biology 5:e1000436.

Chen E Introduction to Latent Dirichlet Allocation. In: http://blog.echen.me/2011/08/22/introduction-to-latent-dirichlet-allocation/.

Crosse MJ, Di Liberto GM, Bednar A, Lalor EC (2016) The Multivariate Temporal Response Function (mTRF) Toolbox: A MATLAB Toolbox for Relating Neural Signals to Continuous Stimuli. Front Hum Neurosci 10:604.

Dale AM, Fischl B, Sereno MI (1999) Cortical surface-based analysis. I. Segmentation and surface reconstruction. NeuroImage 9:179–194.

Dale AM, Liu AK, Fischl BR, Buckner RL, Belliveau JW, Lewine JD, Halgren E (2000) Dynamic statistical parametric mapping: combining fMRI and MEG for high-resolution imaging of cortical activity. Neuron 26:55–67.

de Jong NH, Wempe T (2009) Praat script to detect syllable nuclei and measure speech rate automatically. Behav Res Methods 41:385–390.

Flinker A, Doyle WK, Mehta AD, Devinsky O, Poeppel D (2019) Spectrotemporal modulation provides a unifying framework for auditory cortical asymmetries. Nat Hum Behav 3:393–405.

Gramfort A, Luessi M, Larson E, Engemann DA, Strohmeier D, Brodbeck C, Parkkonen L, Hamalainen MS (2014) MNE software for processing MEG and EEG data. NeuroImage 86:446–460.

Gramfort A, Luessi M, Larson E, Engemann DA, Strohmeier D, Brodbeck C, Goj R, Jas M, Brooks T, Parkkonen L, Hamalainen M (2013) MEG and EEG data analysis with MNE-Python. Front Neurosci 7:267.

Griffiths TL, Steyvers M, Tenenbaum JB (2007) Topics in semantic representation. Psychol Rev 114:211–244.

Haider CL, Suess N, Hauswald A, Park H, Weisz N (2022) Masking of the mouth area impairs reconstruction of acoustic speech features and higher-level segmentational features in the presence of a distractor speaker. NeuroImage 252:119044.

Har-Shai Yahav P, Zion Golumbic E (2021) Linguistic processing of task-irrelevant speech at a cocktail party. Elife 10.

Haufe S, Meinecke F, Gorgen K, Dahne S, Haynes JD, Blankertz B, Biessmann F (2014) On the interpretation of weight vectors of linear models in multivariate neuroimaging. NeuroImage 87:96–110.

Hauswald A, Lithari C, Collignon O, Leonardelli E, Weisz N (2018) A Visual Cortical Network for Deriving Phonological Information from Intelligible Lip Movements. Curr Biol 28:1453–1459 e1453.

Hickok G (2010) The role of mirror neurons in speech and language processing. Brain Lang 112:1–2.

Hoffman P (2019) Reductions in prefrontal activation predict off-topic utterances during speech production. Nat Commun 10:515.

Honnibal M, Montani I (2017) spaCy 2: Natural language understanding with Bloom embeddings, convolutional neural networks and incremental parsing.

Huth AG, de Heer WA, Griffiths TL, Theunissen FE, Gallant JL (2016) Natural speech reveals the semantic maps that tile human cerebral cortex. Nature 532:453–458.

Kaufeld G, Bosker HR, Ten Oever S, Alday PM, Meyer AS, Martin AE (2020) Linguistic Structure and Meaning Organize Neural Oscillations into a Content-Specific Hierarchy. J Neurosci 40:9467–9475.

Kong YY, Mullangi A, Ding N (2014) Differential modulation of auditory responses to attended and unattended speech in different listening conditions. Hear Res 316:73–81.

Koskinen M, Kurimo M, Gross J, Hyvarinen A, Hari R (2020) Brain activity reflects the predictability of word sequences in listened continuous speech. NeuroImage 219:116936.

Kullback S, Leibler RA (1951) On Information and Sufficiency. Ann Math Stat 22:79–86.

Kutas M, Federmeier KD (2011) Thirty years and counting: finding meaning in the N400 component of the event-related brain potential (ERP). Annu Rev Psychol 62:621–647.

Lalor EC, Pearlmutter BA, Reilly RB, McDarby G, Foxe JJ (2006) The VESPA: a method for the rapid estimation of a visual evoked potential. NeuroImage 32:1549–1561.

Landauer TK, Dumais ST (1997) A solution to Plato’s problem: The latent semantic analysis theory of acquisition, induction, and representation of knowledge. Psychological Review 104:211–240.

Likert R (1932) A technique for the measurement of attitudes. Archives of Psychology 22:1–55.

Maris E, Oostenveld R (2007) Nonparametric statistical testing of EEG-and MEG-data. J Neurosci Methods 164:177–190.

MATLAB (R2019b) Natick, Massachusetts. The MathWorks Inc.

Meister IG, Wilson SM, Deblieck C, Wu AD, Iacoboni M (2007) The essential role of premotor cortex in speech perception. Curr Biol 17:1692–1696.

Mikolov T, Chen K, Corrado G, Dean J (2013) Efficient estimation of word representations in vector space. arXiv 1301.3781.

Morillon B, Hackett TA, Kajikawa Y, Schroeder CE (2015) Predictive motor control of sensory dynamics in auditory active sensing. Curr Opin Neurobiol 31:230–238.

Nishida S, Blanc A, Maeda N, Kado M, Nishimoto S (2021) Behavioral correlates of cortical semantic representations modeled by word vectors. PLoS computational biology 17:e1009138.

Nolte G (2003) The magnetic lead field theorem in the quasi-static approximation and its use for magnetoencephalography forward calculation in realistic volume conductors. Physics in medicine and biology 48:3637–3652.

Oldfield RC (1971) The assessment and analysis of handedness: the Edinburgh inventory. Neuropsychologia 9:97–113.

Oostenveld R, Fries P, Maris E, Schoffelen JM (2011) FieldTrip: Open source software for advanced analysis of MEG, EEG, and invasive electrophysiological data. Computational intelligence and neuroscience 2011:156869.

Park H, Thut G, Gross J (2018a) Predictive entrainment of natural speech through two fronto-motor top-down channels. Language, Cognition and Neuroscience:1–13.

Park H, Kayser C, Thut G, Gross J (2016) Lip movements entrain the observers’ low-frequency brain oscillations to facilitate speech intelligibility. Elife 5:e14521.

Park H, Ince RA, Schyns PG, Thut G, Gross J (2015) Frontal top-down signals increase coupling of auditory low-frequency oscillations to continuous speech in human listeners. Curr Biol 25:1649–1653.

Park H, Ince RAA, Schyns PG, Thut G, Gross J (2018b) Representational interactions during audiovisual speech entrainment: Redundancy in left posterior superior temporal gyrus and synergy in left motor cortex. PLoS biology 16:e2006558.

Pedregosa F, Varoquaux G, Gramfort A, Michel V, Thirion B, Grisel O, Blondel M, Prettenhofer P, Weiss R, Dubourg V, Vanderplas J, Passos A, Cournapeau D, Brucher M, Perrot M, Duchesnay E (2011) Scikit-learn: Machine Learning in Python. J Mach Learn Res 12:2825–2830.

Pereira F, Detre G, Botvinick M (2011) Generating text from functional brain images. Front Hum Neurosci 5:72.

Pereira F, Lou B, Pritchett B, Ritter S, Gershman SJ, Kanwisher N, Botvinick M, Fedorenko E (2018) Toward a universal decoder of linguistic meaning from brain activation. Nat Commun 9:963.

Skipper JI, Devlin JT, Lametti DR (2017) The hearing ear is always found close to the speaking tongue: Review of the role of the motor system in speech perception. Brain and Language 164:77–105.

Smith ZM, Delgutte B, Oxenham AJ (2002) Chimaeric sounds reveal dichotomies in auditory perception. Nature 416:87–90.

Strauss A, Kotz SA, Scharinger M, Obleser J (2014) Alpha and theta brain oscillations index dissociable processes in spoken word recognition. NeuroImage 97:387–395.

Team RC (2020) R: A language and environment for statistical computing. R Foundation for Statistical Computing.

Teng X, Meng Q, Poeppel D (2021) Modulation Spectra Capture EEG Responses to Speech Signals and Drive Distinct Temporal Response Functions. eNeuro 8.

Vallat R (2018) Pingouin: statistics in Python. Journal of Open Source Software 3:1026.

van der Maaten LJP, Hinton GE (2008) Visualizing Data using t-SNE. J Mach Learn Res 9:2579–2605.

Van Essen DC (2005) A Population-Average, Landmark- and Surface-based (PALS) atlas of human cerebral cortex. NeuroImage 28:635–662.

Wang L, Kuperberg G, Jensen O (2018) Specific lexico-semantic predictions are associated with unique spatial and temporal patterns of neural activity. Elife 7.

Weischedel R, Palmer M, Marcus M, Hovy E, Pradhan S, Ramshaw L, Xue N, Taylor A, Kaufman J, Franchini M, El-Bachouti M, Belvin R, Houston A (2013) OntoNotes Release 5.0 LDC2013T19. Web Download. Philadelphia: Linguistic Data Consortium.

Wickham H (2016) ggplot2: Elegant Graphics for Data Analysis: Springer-Verlag New York.

Yu G (2020) scatterpie: Scatter Pie Plot. In, R package version 0.1.5. Edition.

