## Supplementary Information for "Get the gist of the story: Neural map of topic keywords in multi-speaker environment"

**Table S1 (related to Fig. 1a). Profile of each talk used in the study**

| Talk | Duration (s) | Phonation Time (s) | Number of Syllables | Speech Rate | Articulation Rate | Number of Chunks |
|---|---|---|---|---|---|---|
| Talk 1 | 432.45 | 327.55 | 1420 | 3.28 | 4.34 | 145 |
| Talk 2 | 461.41 | 404.82 | 1671 | 3.62 | 4.13 | 108 |
| Talk 3 | 389.98 | 351.89 | 1382 | 3.54 | 3.93 | 95 |
| Talk 4 | 542.77 | 463.46 | 1911 | 3.52 | 4.12 | 135 |
| Talk 5 | 536.43 | 460.38 | 1912 | 3.56 | 4.15 | 144 |
| Talk 6 | 584.72 | 526.34 | 2167 | 3.71 | 4.12 | 138 |
| Talk 7 | 513.04 | 444.43 | 1857 | 3.62 | 4.18 | 140 |
| **Mean** | 494.40 | 425.55 | 1760.00 | 3.55 | 4.14 | 129.29 |
| **SD** | 68.85 | 69.07 | 285.01 | 0.13 | 0.12 | 19.64 |

Using the Syllable Nuclei library (de Jong and Wempe, 2009) in Praat (Boersma and Weenink, 2018), we derived acoustically segmented speech chunks for topic modelling analysis (see Fig. 1a). For detecting acoustic silences, we used parameters of 0.25 s-long minimum pause (silence) duration and 25 dB silence threshold. The duration of each speech chunk was 1 s-long at the minimum. One out of eight talks (see Stimuli and Experiment in Materials and Methods) was used for visual speech only for other experimental conditions, so we analyzed seven talks only. The table shows the number of syllables, speech rate (the number of syllables divided by the duration of the talk), articulation rate (the number of syllables divided by phonation (speaking) time), and the number of segmented speech chunks for each talk.

**Sentiment analysis**

The selected TED talks are all taken from informative, persuasive, inspiring categories; however, in order to further rule out the possibility of the neural representation of topic keywords processing driven by valence, we performed sentiment analysis on speech materials.

To discern sentiment on the raw text, we used VADER (Valence Aware Dictionary for sEntiment Reasoning) (Hutto and Gilbert, 2014) model in the NLTK 3.5 (Natural Language Toolkit) package (Bird et al., 2009), which provides both polarities (positive/neutral/negative) and intensity (strength) of emotion. For instance, a word like “good” or “awesome” is labelled as positive valence, whereas “adverse” or “betrayal” is labelled as negative valence. VADER model relies on a dictionary that maps lexical features to sentiment scores (emotion intensities), and the total sentiment score of a text is obtained by summing up the intensity of each word in the text. This model is able to discern “love” as positive and “did not love” as negative sentiment. Also, the model adjusts the score based on the intensity of a word, for example, a boosted score for “totally” (+0.293) and a reduced score for “slightly” (-0.293), as well as for capitalization and punctuation (e.g., more scores for “GREAT” than “great”). The limitation of the sentiment analysis in general lies in its discernibility of sarcastic expression using positive words in a negative way. The model returns scores for each category of positive, neutral, negative and compound. The maximum values for positive, neutral and negative are 1.0, and the compound score is a normalized value across positive, neutral, negative scores to be between -1 and 1 using an alpha ($\alpha$=15) that approximates the max expected value. Compound scores above, near and below zero indicate positive, neutral and negative valence, respectively.

In the current study, we performed the sentiment analysis on each segmented speech chunk resulting in a score for each speech chunk in a talk. Then the compound scores were averaged across speech chunks in a talk, as shown in Supplementary Table 2. The results of near-zero compound scores prove that the speech materials used in the current study are all with a neutral sentiment (also shown in the high scores for neutral sentiment; see the table below), so they presumably have no effects on the attentional shift between attended (to-be-attended) and unattended (to-be-ignored) talks due to emotional contents or stronger high-level semantic processing for one talk over another.

**Table S2. Sentiment analysis for each talk used in the study**

| | | Talk 1 | Talk 2 | Talk 3 | Talk 4 | Talk 5 | Talk 6 | Talk 7 |
|---|---|---|---|---|---|---|---|---|
| Compound | mean | **0.12** | **0.10** | **0.12** | **0.16** | **0.04** | **0.05** | **0.01** |
| | std | 0.27 | 0.35 | 0.26 | 0.28 | 0.25 | 0.28 | 0.33 |
| | min | -0.75 | -0.88 | -0.54 | -0.67 | -0.69 | -0.73 | -0.79 |
| | max | 0.82 | 0.95 | 0.82 | 0.88 | 0.88 | 0.86 | 0.91 |
| Neutral | mean | **0.85** | **0.85** | **0.88** | **0.82** | **0.90** | **0.90** | **0.84** |
| | std | 0.20 | 0.19 | 0.16 | 0.20 | 0.15 | 0.16 | 0.19 |
| | min | 0.31 | 0.21 | 0.15 | 0.18 | 0.39 | 0.36 | 0.21 |
| | max | 1.00 | 1.00 | 1.00 | 1.00 | 1.00 | 1.00 | 1.00 |
| Positive | mean | 0.12 | 0.11 | 0.09 | 0.14 | 0.06 | 0.07 | 0.08 |
| | std | 0.18 | 0.16 | 0.13 | 0.17 | 0.13 | 0.13 | 0.15 |
| | min | 0.00 | 0.00 | 0.00 | 0.00 | 0.00 | 0.00 | 0.00 |
| | max | 0.68 | 0.69 | 0.55 | 0.59 | 0.62 | 0.57 | 0.79 |
| Negative | mean | 0.04 | 0.04 | 0.03 | 0.04 | 0.04 | 0.04 | 0.08 |
| | std | 0.10 | 0.13 | 0.10 | 0.10 | 0.09 | 0.11 | 0.14 |
| | min | 0.00 | 0.00 | 0.00 | 0.00 | 0.00 | 0.00 | 0.00 |
| | max | 0.52 | 0.79 | 0.55 | 0.52 | 0.51 | 0.64 | 0.49 |

**Spatio-temporal clusters of encoding neural map for topic keywords processing**

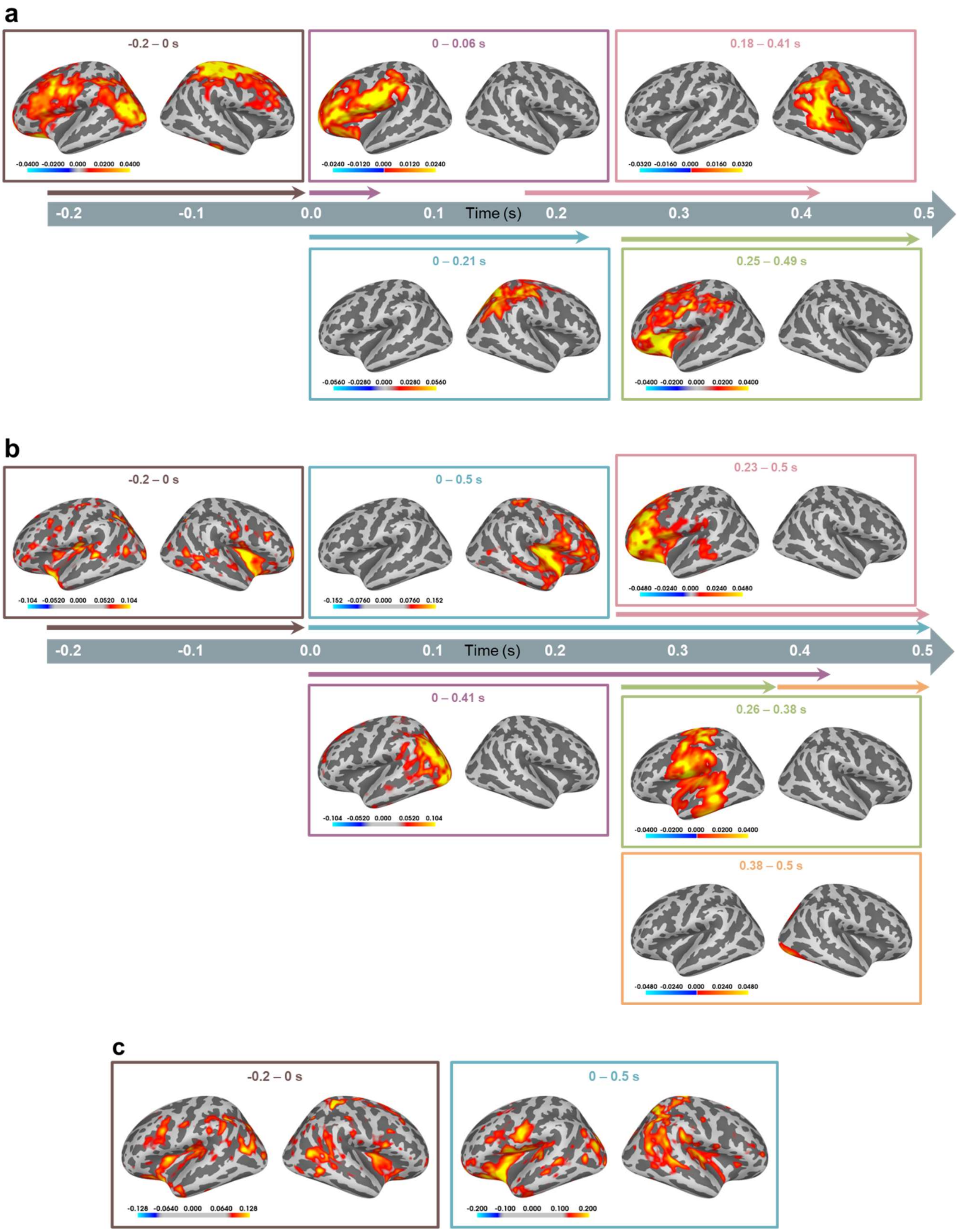

**Supplementary Figure 1 (related to Fig. 4). Topic keywords neural map: Spatio-temporal clusters of difference between high vs. low topic probability conditions using encoding model coefficients**. **a,** (related to Fig. 4a) Attended, High vs. Attended, Low. **b,** (related to Fig. 4b) Unattended, High vs. Unattended, Low. **c,** (related to Fig. 4c) Unattended, High vs. Attended, Low. Colorbars indicate scaled t-values computed using two-tailed cluster-level spatio-temporal permutation t-test ($p < 0.05$, 1024 permutations). T-values in significant clusters are scaled corresponding to the duration spanned by the cluster (for more details, see Statistical test in Materials and Methods).

We investigated the difference between high and low topic probability derived from both attended and unattended talks for 0 - 0.5 s with respect to the onset of speech chunk using encoding model coefficients (Fig. 4). The sensor-level model coefficients were mapped onto the brain surface using the dynamic statistical parametric mapping (dSPM) source localization method. The spatio-temporal difference between high vs. low topic probability was computed via a cluster-level spatio-temporal permutation t-test ($p < 0.05$), and here we show the map for before speech onset (-0.2 and 0 s) as well in order to investigate the predictive neural processing before the onset of speech chunks (i.e., pauses between articulations) in connected speech. Detailed temporal information of each spatial-temporal cluster is shown as a timeline.

**a,** (related to Fig. 4a) Attended, High vs. Attended, Low. After speech onset, the activities of the model contribution are stronger in the left inferior frontal (BA 44, 45), somatosensory areas, as well as primary motor (BA 4), premotor (BA 6) cortices (0 to 0.06 s) and right parietal/temporal areas, including temporo-parieto-occipital junction (0 s to 0.21 s) for high compared to low topic probability speech in attended talk. Then, significant differences emerge in the right inferior parietal and temporal cortex between 0.18 s to 0.41 s. During the 0.25 s to 0.49 period, the difference is located in the left inferior frontal, motor and opercular areas. Before speech onset, the left inferior temporal, premotor/motor cortex and visual cortex and right somatosensory and premotor cortex are shown to be stronger for speech chunks with high topic probability compared to low topic probability.

**b,** (related to Fig. 4b) Unattended, High vs. Unattended, Low. After speech onset, the differences are localized in the right frontal and temporal areas, left frontal, posterior temporal and visual cortex stronger for high compared to low topic probability for the entire analysis window (0 s to 0.5 s). Left inferior parietal and lateral occipital cortices are also stronger for the high than the low topic probability for ~0.4 s (0 s to 0.41 s). Then the difference is shown in the left inferior frontal (BA 45, 47) and dorsolateral prefrontal (BA 9, 46) areas (0.23 s to 0.5 s). During the similar but shorter time interval (0.26 s to 0.38 s), left somatosensory including motor, premotor areas, primary auditory and auditory association cortices including superior temporal gyrus/sulcus are shown significantly stronger for high vs. low topic probability. Before speech onset, right inferior frontal areas are shown stronger for speech chunks with high topic probability compared to low topic probability.

**c,** (related to Fig. 4c) Unattended, High vs. Attended, Low. For both before and after the speech onset duration, the difference was not observed as subsets of temporal clusters. Thus, the neural map for after the speech onset (0 - 0.5 s) is the same as in Fig. 4c. The neural map for before the speech onset (-0.2 - 0 s) shows a similar spatial pattern but to a lesser extent as the neural map for after the speech onset.

**Decoding neural map for topic keywords processing**

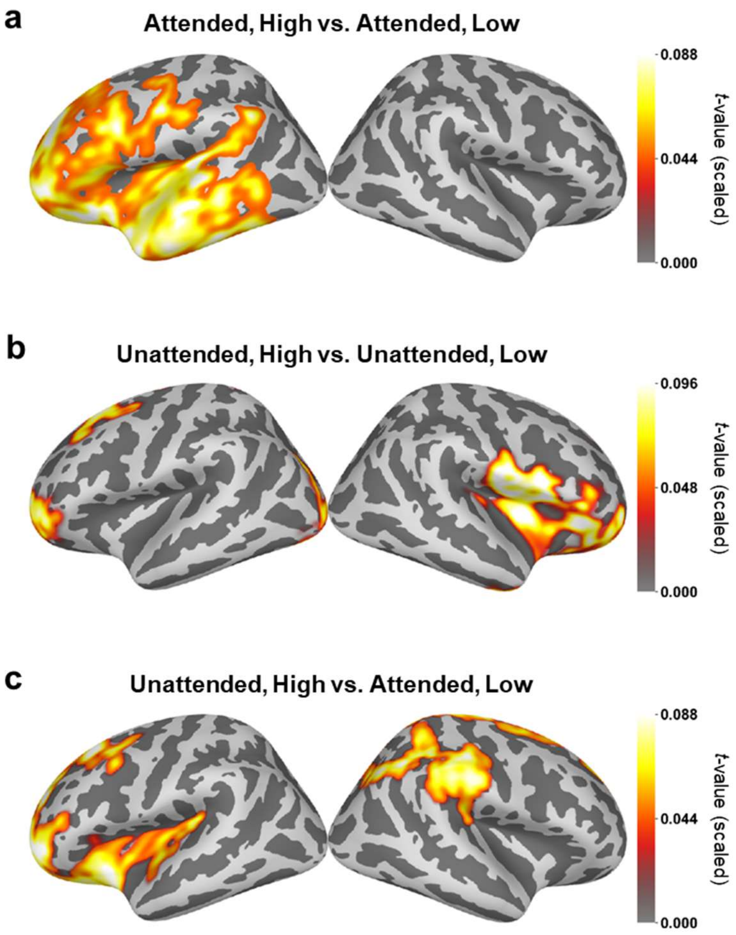


**Supplementary Figure 2 (related to Fig. 4). Topic keywords neural map from inverse-transformed decoding model coefficients.**

Neural entrainment to complex, time-varying speech stimuli can be modelled via multivariate temporal response function (mTRF). We used mTRF to investigate the difference between the processing of speech with high vs. low topic probability and showed the encoding results in Fig. 4 and Supplementary Figure 1. The decoding (backward: brain to stimulus, stimulus reconstruction) model offers a complementary perspective on this mapping by reconstructing stimulus features (i.e., speech envelope). Neurophysiological interpretation for the decoding model results has been known to be plausible via inverse transformed decoder model weights. Decoding model weights are not interpretable in a neurophysiological sense because significant nonzero weights could be observed, which are statistically independent of the neural responses; however, transforming the backward model into the forward model enables neurophysiological interpretation of the backward model weights (Haufe et al.,

2014). We applied this inverse procedure to transform decoder weights to forward model space on brain source space via the dSPM method.

In the decoding model, each of the MEG sensors is treated as an independent feature in a multivariate context. As shown in the previous literature (Gross et al., 2013; Park et al., 2016), given the relationship between the speech and neural responses by ~100 ms conduction delay, here we set negative time delay by ~100 ms for MEG data (-0.1 s to 0 s in steps of 8 ms). The decoder is then computed using linear regression (least-squares loss function), and reconstructed speech and original speech are compared by the model performance metric, the correlation coefficient (prediction score, *r*; Fig. 3b). The K-fold (k=3) cross-validation was used to split data into train and test sets (5:5), and fitting the model was iterated through train-test sets. During the model fitting, decoder model coefficients (filter weights) and inverse-transformed coefficients (also known as forward mixing weights) (Haufe et al., 2014) were obtained (number of cross-validation x number of sensors x time delays: 3 x 248 x 13). Model coefficients and inverse-transformed coefficients are averaged across cross-validation iteration and time delays. Inverse-transformed coefficients map provides decoded patterns that explain how the measured data was generated from the discriminant neural sources extracted by the filter weights, thus neurophysiologically more meaningful and interpretable than the coefficients map. This decoded patterns map was further investigated via dSPM source inverse solution. The computation was performed separately for each condition (high and low topic probability from each attended and unattended talk), and the statistical significance map between the conditions was derived using two-tailed cluster-level spatio-temporal permutation t-test ($p < 0.05$, 1024 permutations). T-values in significant clusters are scaled corresponding to the duration spanned by the cluster (for more details, see Statistical test in Materials and Methods).

**a,** (related to Fig. 4a) Attended, High vs. Attended, Low. The decoded patterns map is localized in the left inferior and dorsolateral frontal, motor areas as well as extensive auditory and temporal cortices. **b,** (related to Fig. 4b) Unattended, High vs. Unattended, Low. The decoded patterns map is localized in the right inferior frontal/insular areas, left superior/dorsolateral prefrontal areas and bilateral anterior/middle cingulate cortices. **c,** (related to Fig. 4c) Unattended, High vs. Attended, Low. The decoded patterns map is localized in the left inferior frontal and opercular areas and dorsolateral prefrontal cortices and the right parietal area.

Brain areas found in the encoding map and decoded patterns maps show quite an extensive overlap, albeit exhibiting differences.

**Encoding model contribution in each mediator region for all conditions**

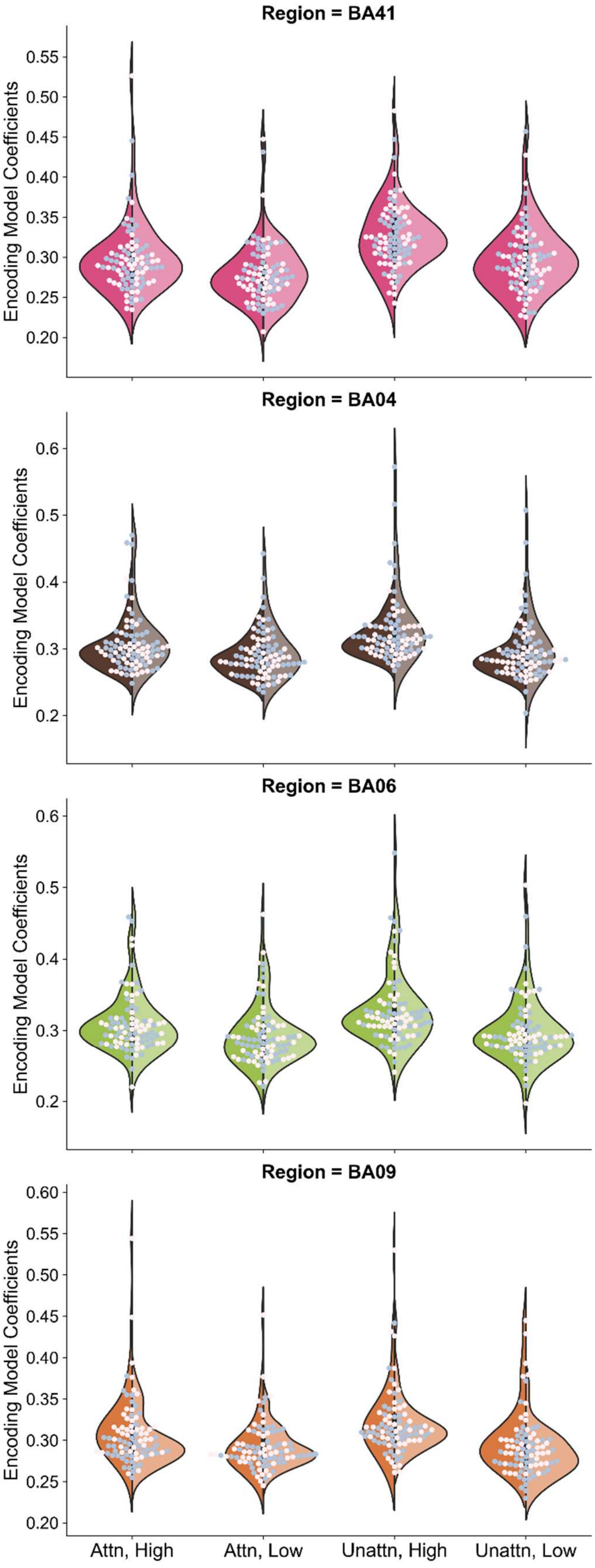

**Supplementary Figure 3 (related to Fig. 5). Encoding model coefficients in mediator regions of the PALS-B12-Brodmann atlas.**

Here we provide more extensive results in relation to the mediation results shown in Figure 5. The encoding model coefficients in each condition from the combinations of attention and topic probability are displayed for individual mediator brain regions (Fig. 5a) in the parcellations of the PALS-B12-Brodmann atlas as subplots. In each subplot, x- and y-axes depict conditions (attended high, attended low, unattended high, unattended low) and encoding model coefficients, respectively. In each violin plot, the left and right side distribution represent a kernel density estimation from the left and right hemispheres of the region. Each dot represents each subject's data in light pink for the left hemisphere and in light gray for the right hemisphere.

### SI References


Bird S, Ewan K, Loper E (2009) Natural Language Processing with Python: O'Reilly Media, Inc.

Boersma P, Weenink D (2018) Praat: doing phonetics by computer [Computer program]. Version 6.0.37:retrieved 14 March 2018 from http://www.praat.org/.

de Jong NH, Wempe T (2009) Praat script to detect syllable nuclei and measure speech rate automatically. Behav Res Methods 41:385-390.

Gross J, Hoogenboom N, Thut G, Schyns P, Panzeri S, Belin P, Garrod S (2013) Speech rhythms and multiplexed oscillatory sensory coding in the human brain. PLoS biology 11:e1001752.

Haufe S, Meinecke F, Gorgen K, Dahne S, Haynes JD, Blankertz B, Biessmann F (2014) On the interpretation of weight vectors of linear models in multivariate neuroimaging. NeuroImage 87:96-110.

Hutto CJ, Gilbert EE (2014) VADER: A Parsimonious Rule-based Model for Sentiment Analysis of Social Media Text. In: Eighth International Conference on Weblogs and Social Media (ICWSM-14). Ann Arbor, MI.

Park H, Kayser C, Thut G, Gross J (2016) Lip movements entrain the observers' low-frequency brain oscillations to facilitate speech intelligibility. Elife 5:e14521.
